# Modified covariance beamformer for solving MEG inverse problem in the environment with correlated sources

**DOI:** 10.1101/668814

**Authors:** Aleksandra Kuznetsova, Yulia Nurislamova, Alexei Ossadtchi

## Abstract

Magnetoencephalography (MEG) is a neuroimaging method ideally suited for non-invasive studies of brain dynamics. MEG’s spatial resolution critically depends on the approach used to solve the ill-posed inverse problem in order to transform sensor signals into cortical activation maps. Over recent years non-globally optimized solutions based on the use of adaptive beamformers (BF) gained popularity.

When operating in the environment with a small number of uncorrelated sources the BFs perform optimally and yield spatial super-resolution. However, the BFs are known to fail when dealing with correlated sources acting like poorly tuned spatial filters with low signal-to-noise ratio (SNR) of the output timeseries and often meaningless cortical maps of power distribution.

This fact poses a serious limitation on the broader use of this promising technique especially since fundamental mechanisms of brain functioning, its inherent symmetry and task-based experimental paradigms result into a great deal of correlation in the activity of cortical sources. To cope with this problem, we developed a novel beamformer approach that preserves high spatial resolution in the environments with correlated sources.

At the core of our method is a projection operation applied to the vectorized sensor-space covariance matrix. This projection does not remove the activity of the correlated sources from the sensor-space covariance matrix but rather selectively handles their contributions to the covariance matrix and creates a sufficiently accurate approximation of an ideal data covariance that could hypothetically be observed should these sources be uncorrelated. Since the projection operation is *reciprocal* to the PSIICOS method developed by us earlier (Ossadtchi et al. (2018)) we refer to the family of algorithms presented here as ReciPSIICOS.

We asses the performance of the novel approach using realistically simulated MEG data and show its superior performance in comparison to the well established MNE and classical BF approaches. We have also applied our approach to the MEG datasets from the two experiments involving two different auditory tasks.

The analysis of experimental MEG datasets showed that beamformers from ReciPSIICOS family, but not MNE and the classical BF, discovered the expected bilateral focal sources in the primary auditory cortex and detected motor cortex activity associated with the audio-motor task. Moreover, ReciPSIICOS beamformers yielded cortical activity estimates with amplitude an order of magnitude higher than that obtained with the classical BF, which indicates the severeness of the signal cancellation problem when applying classical beamformers to MEG signals generated by synchronous sources.

## 1 Introduction

Magnetoencephalography (MEG) is a noninvasive neuroimaging method hallmarked by the millisecond scale temporal resolution and subcentimeter spatial resolution. As such, this method is very well suited for studying the fine features of spatio-temporal dynamics exhibited by neural circuits. The high temporal resolution of MEG is concomitant to the nature of the underlying electrophysiological processes in the brain tissue. As to the spatial resolution, it crucially depends on how the ill-posed inverse problem is approached to recover the distribution of neural sources from the magnetic field measurements around the head.

Among the plethora of methods for solving the MEG inverse problem (Hämäläinen and Ilmoniemi (1994), Pascual-Marqui et al. (1994), Gorodnitsky et al. (1995), Matsuura and Okabe (1995), Mosher and Leahy (1999)) local linear estimators such as adaptive linearly constrained minimum variance (LCMV) beamformers (Van Veen et al. (1997), Sekihara et al. (2001), Greenblatt et al. (2005)) are known for their high spatial resolution that can be achieved under the proper conditions, namely when the measurements are generated by a small number of focal sources, and these sources are uncorrelated.

In reality, these assumptions are often violated. Thus, cortical sources exhibit transient synchrony (Varela et al. (2001)) that manifests ongoing integrative processes (Fries (2015)). One reason for such synchronization is the time-locking of brain activity (called event-related potentials, ERPs) to task events, such as movement or stimulus onset, for a wide range of cognitive, motor and sensory paradigms (Gascoyne et al. (2016)). Synchronous ERPs often occur in both hemispheres at the functionally homologous areas. Correlated sources cause a significant reduction in SNR when adaptive beamformers are used to process the data (Sekihara and Nagarajan (2008)).

Several approaches have been developed to improve beamforming in the presence of correlated sources. Dalal et al. (2006) suggested that the entire region that may potentially include a source correlated to the activity in the region of interest (ROI) should be suppressed. This idea can be implemented using an SVD derived constraint based on the topography of the cortical patch containing the interfering source. This approach requires an *apriori* knowledge of the locations of correlated sources. Nulling the activity of multiple regions (or spatially-extended ones) with this method reduces the number of degrees of freedom, that otherwise could be used for suppressing the interfering sources.

Brookes et al. (2007) suggested building a beamformer based on the constraints that are calculated from the topographies of correlated sources using their linear combination. An “amplitude optimization” routine was suggested to compute the optimal mixing coefficients for this procedure. Application of this method to real data requires explicit scanning over all possible pairs using a coarse grid, which is not time efficient and prone to errors if the seed source is set *apriori*.

Two beamformers that allow to overcome correlated sources issue were evaluated in Popescu et al. (2008): (1) a linearly constrained minimum variance beamformer with partial sensor coverage (LCMV-PSC), and (2) a multiple constrained minimum-variance beamformer with coherent source region suppression (MCMV-CSRS). It was demonstrated that the latter exhibits precise localization and minimal amplitude and phase distortion for a broad range of relative positions of the interfering source within the suppression region. With this method, again, degrees of freedom are consumed because of the assumption regarding the location of the interfering source that maintains the regional zero constraint.

Quraan and Cheyne (2010) compared various solutions available to the date of that publication to cope with correlated sources in beamforming. When prior information about the location of correlated sources is available, the method of beamforming with coherent source region suppression described in Dalal et al. (2006) appeared to be the most effective, including the case of closely located (3 cm apart) correlated sources. The authors concluded that this solution, when carefully exercised, can significantly improve localization accuracy, but does not fully solve the amplitude bias problem.

Diwakar et al. (2011) introduced a dual-core beamforming idea, which is an extension of Sekihara’s vectorized LCMV approach (Sekihara et al. (2001)). Dual-core beamformer is built using the constraint created from the topographies of two spatially disjoint regions. This approach allows to find the pairs of highly correlated sources and eliminate the computationally expensive search for topographies mixing parameter and optimal dipole orientation required in the approach of Brookes et al. (2007).

The approaches based on finding pairs of correlated sources would, for example, fail to detect a hub coupled simultaneously to more than one additional source. Further, as our simulations show, synchrony between more than two sources has a complex effect on the suppression of the beamformer output power. Therefore, in a number of practical situation, the beamformers limited to considering only a pair of sources could fail. Moiseev et al. (2011) presented a detailed treatment of the multiple constrained minimum-variance beamformers with coherent source region suppression (MCMV-CSRS) and offered a set of practical solutions and scanning statistics to be used for identifying cortical regions with correlated activity.

The approaches for coping with source synchrony problem that we have described so far are conceptually similar and spin around the idea of suppressing the activity of the sources correlated to the current ROI. Additional insights into the problem of source correlation could be gained from the data covariance matrix, which contains the information about source synchrony. Since beamforming algorithms are based on the premise of the absence of correlation of the underlying sources, the covariance matrix may get incorrectly interpreted by the adaptive beamforming algorithm when this premise is violated. The beamformer weights calculated using such a covariance matrix would result in a significant suppression of the SNR of the estimated source timeseries, and in the case of perfect synchrony the beamformer would completely cancel the signal.

The approach we propose here is based on the analysis of data covariance matrix. Kimura et al. (2007) previously developed a method that is conceptually close to ours. They used a forward model and the least squares approximation approach to find an estimate of the source-space covariance matrix that corresponded to the interaction of a small number of sources. They then nulled the off-diagonal elements of this matrix and projected it back to the sensor space. This new decorrelated matrix was used for building the conventional beamformers. The super-resolution properties of this approach depend on the estimated number of active sources chosen to comprise the source-space covariance matrix, as well as on the other parameters of the procedure employed to find the least squares approximation of the source-space covariance matrix. The selection of the number of sources to be represented in the source-space covariance matrix is a non-trivial problem that affects the performance of this technique.

In this paper, we propose a way to modify data covariance matrix that makes adaptive LCMV beamforming robust against correlation of neural sources activity. The proposed procedure selectively handles the data covariance components and creates a close approximation of an ideal data covariance that could hypothetically be observed in the absence of correlation of source timeseries. In contrast to the approach described in Kimura et al. (2007), our procedure is data-independent. It relies only on the forward model and can be considered as a deterministic extra step in building an adaptive LCMV-based inverse operator. By design, our procedure does not attempt to estimate active sources, but rather exploits the spatial structure of the inherent differences between the auto-terms and cross-terms of the covariance matrix.

In what follows, we start with a model of the averaged evoked response data and derive the generative model for the sensor-space covariance matrix. Next, we introduce the basic adaptive beamforming mathematics and illustrate that the LCMV beafmromer is merely a match filter operating in the space of the whitened data with respect to the data covariance matrix. We then analytically explore the effects of non-zero correlation between source timeseries in the environment with three sources and show that not only the activity of a pair of strongly correlated sources gets significantly suppressed when estimated with adaptive beamformer but also the third source with relatively weak coupling to the activity of the first pair of sources suffers a drastic reduction of amplitude when estimated by the beamformer. This motivates us to develop a procedure that does not rely on the potentially erroneous source estimation step.

Following these steps, we return to the analytic expression for the data covariance matrix and show that its vectorized version can be considered as a linear combination of the components modulated solely by the power of sources and those modulated by the off-diagonal elements of the source-space covariance matrix. Based on this observation, we propose two simple projection procedures to suppress the components of the data covariance matrix that are modulated by the synchrony of neuronal sources. Next we employ simulations to explore the performance of our method under various conditions and compare it against the MNE solution and adaptive LCMV beamformer. Finally, we apply these techniques to two experimental MEG datasets, one from an audio-motor task and the other from a passive tone-listening task. Finally, we discuss the significance of our results, outline the advantages of the proposed method, identify the potential shortcomings of our approach, and sketch the future directions for the development of the proposed algorithm.

## 2 Methods

### 2.1 Data model

Event-related potentials (ERPs) measured with electrophysiological methods such as electroencephalography (EEG) and magnetoencephalography (MEG) can be sufficiently accurately modelled as a superposition of the contributions from a finite number of sources. Only the contributions that are sufficiently phase-locked to the task onset moment make it through the averaging procedure and form the evoked response (Luck and Kappenman (2011)).

The observation equation linking vector 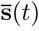 of the averaged source-space activity with vector measured by the 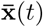 array of *M* sensors at time instance *t* can be written as

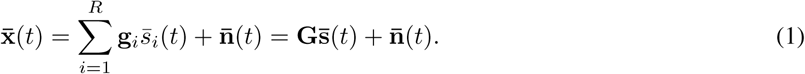

where 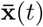 is [*M* × 1] vector and 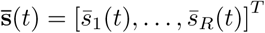 is an [*R* × 1] vector, as we assume, that this evoked activity is generated by a relatively small number *R* of focal cortical sources. **G** = [**g**_1_, …, **g**_*R*_] is a matrix of corresponding source topographies. Noise term 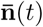 represents the sum of the remainders of the induced and task unrelated activity that is supposed to be sufficiently suppressed by the event-related averaging procedure. One of the key assumptions implicit in this ERP data model is that the stimulus-locked cortical activation profiles are expected to be focal. The number of active stimulus-related sources *R* is usually unknown, but it is assumed to be several orders of magnitude less than the total number of cortical sources.

The activity is observed over a finite period of *T* samples. Where appropriate, we will use the following matrix formulation of (1)

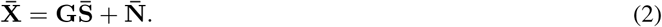

Here 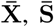 and 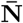 are the matrices comprising sensor recordings 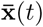, source activation profiles 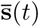 and noise measurements 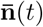 at consecutive time instances *t* = 1, …, *T*.

The active source locations are unknown, and finding them is the goal of the EEG and MEG inverse problem solving. We approach this daunting problem with the knowledge of the forward model that matches every *i*-th location of the dipolar sources with a topography vector **g**_*i*_ that contains the weights for the contribution of the *i*-th unit dipole to the sensor measurements. Our goal is then to identify the grid nodes containing active dipoles. Sufficient accuracy of computing **g**_*i*_ is a strong requirement to ensure adequate performance of the inverse solvers, including adaptive beamformers.

Within the adaptive beamforming approach, when the forward model is not accurate enough the model-based and actual topographies of the source at a given location do not match. In this case, the adaptive beamformers would exhibit signal cancellation phenomenon manifested as a significant reduction of the signal-to-noise ratio (SNR) in the reconstructed source timeseries. This deficiency causes significant source localization errors when adaptive beamformers are used in the scanning mode (Steinsträter et al. (2010)).

Spatial super-resolution is one of the most attractive features of the adaptive beamforming technique. It is achieved using the data covariance matrix that conveys the information about the subset of active sources to the beamformer. This information is then used by the beamformers to efficiently distribute the available degrees of freedom to suppress these active sources.

### 2.2 Adaptive LCMV beamformer

Adaptive linearly constrained minimum variance (LCMV) beamformer (Van Veen et al. (1997); Sekihara et al. (2001)) is a local linear estimator with a unit gain weight constraint (Greenblatt et al. (2005)). Over the recent years, this approach gained popularity as an efficient inverse solver of the MEG inverse problem (Darvas et al. (2004)).

Due to the fundamental limitations of the electromagnetic inverse problem, it is impossible to globally suppress contribution from all non-target sources. Therefore, within the beamforming approach, the problem of finding the spatial filter weights is formulated locally as minimizing the spatial filter output variance under the unit gain constraint with respect to the source of interest.

Beamformers can be used to estimate the activity of a specific ROI or in a scanning mode to assess the distribution of activity over the entire cortex. Various forms of beamformers exist that can be classified based on the source-space and sensor-space norms (Greenblatt et al. (2005)). Unlike global estimators (MNE, wMNE, MCE, etc.), beamformers tuned to different cortical locations do not depend on each other and their summed output projected back to the source space is generally not supposed to be equal to the measured data **X**.

#### 2.2.1 Basic beamforming mathematics

Consider an elementary cortical dipolar source with free orientation within the locally tangential plane located at **r**_*i*_ = [*x*_*i*_, *y*_*i*_, *z*_*i*_]. To reconstruct its activity via the LCMV beamformer from the MEG data, one uses spatial filter **w**_*i*_ = **w**(**r**_*i*_) calculated as the solution of the following optimization problem (Sekihara et al. (2001))

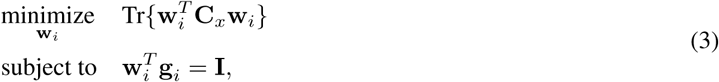

where Tr{·} is a matrix trace operator, (·)^*T*^ is the transpose operator, **C**_*x*_ is the sensor-space covariance matrix, matrix **g**_*i*_ = **G**(**r**_*i*_) contains topographies of the target source and **I** is the identity matrix. With the Lagrange multipliers method, it can be shown that the optimal solution is

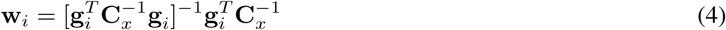

The calculated spatial filters **w** _*i*_ could then be used to reconstruct the source timeseries as

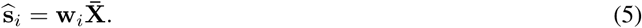

It turns out that the estimated source power 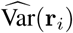 can be computed without the explicit computation of **w**_*i*_:

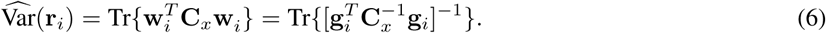

It is also possible to employ the beamformer in the scanning mode and compute power distribution profile 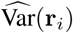 for the entire set of cortical locations **r**_*i*_, *i* = 1, …, *N*. Then, source localization can be performed by searching for the local maximums of the estimated cortical power distribution.

The described approach does not introduce any assumptions on the number of active sources or their spatial distribution. However, the LCMV adaptive beamformer enables spatial super-resolution only in the case when the measured neural activity is produced by a small number of focal cortical sources (Borgiotti and Kaplan (1979)). It is noteworthy that the adaptive LCMV beamforming is technically a biased location estimator Greenblatt et al. (2005).

LCMV beamformer operation principle can be understood if we rewrite (4) using vector 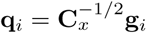 and assume that our target source has fixed orientation and thus **g**_*i*_ is simply an *M* × 1 vector. Then, the timeseries estimated by the beamformer can be written as

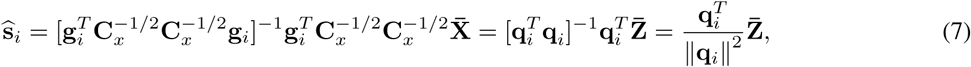

where 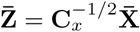 is spatially whitened data. Thus, the beamformer simply whitens the data with respect to the full data covariance matrix and then performs spatial match filtering in this whitened space using the spatial filter equal to the appropriately transformed and normalized target source topography **g**_*i*_. The whitening operation efficiently spreads out all the data variance over the entire data space so that only a small fraction of it leaks into the target source estimate.

#### 2.2.2 Adaptive beamforming in the environment with correlated sources

Despite the high localization efficiency and spatial super-resolution, adaptive LCMV beamformers suffer from the signal cancellation problem when operating in the suboptimal environment. The optimal spatial filters calculated in (4) reflect activity of intrinsic sources correctly only when the measurements are generated by a small number of focal sources with uncorrelated timeseries.

These assumptions, however, are rarely not met for the real experimental conditions because the brain integrative mechanisms result in a high level of synchrony across different areas (Fries (2015)). Thus, synchronous, stimulus-locked responses occur in many cortical areas, such as ERPs occurring in bilateral, functionally homologous locations (Gascoyne et al. (2016)).

Correlation of sources causes significant reduction of the SNR estimated with adaptive beamformers (Sekihara and Nagarajan (2008)). We will start by presenting an analytical demonstration of this phenomenon, first for two sources and then for three sources.

#### 2.2.3 Example of two interacting sources

Consider an example of two active dipolar sources **q**_1_, **q**_2_ with fixed orientations located at points **r**_1_, **r**_2_ and having activation timeseries **s**_1_(*t*), **s**_2_(*t*), *t* = 1 …*T* and source topographies **g**_1_, **g**_2_, correspondingly. The evoked response signal measured with *M* electrodes at *T* time samples can be represented as the following sum: 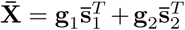. The sensor-space covariance matrix can be rewritten in terms of the source-space covariance matrix elements 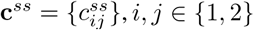:

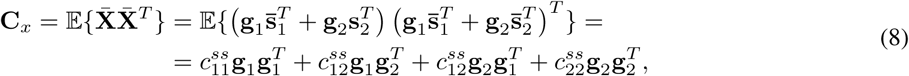

where 𝔼{·} is the expected value operator.

As follows from equations (4) and (6), given the forward model, the covariance matrix fully determines the beamformer weights and the output power of source estimates when applied to the data 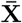.

The true variance of source **q**_1_ is 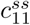 and its LCMV based estimate equals to the value of target function (3) at the optimum (4):

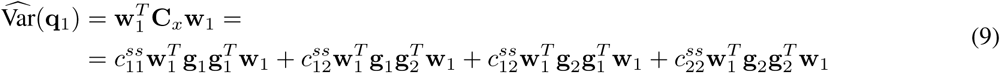

It is straightforward but tedious to show (Sekihara and Nagarajan (2008)), that the estimated power of each source decreases as a quadratic function of the correlation coefficient 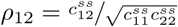, and in the case of unit source variance, the power can be expressed as

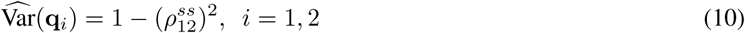

In the case of complete synchrony, the adaptive beamformer will simply output zero. Intuitively this can be understood as follows. In order to meet the constraint, the beamformer provides unit gain for the target source. The functional optimized by the adaptive beamformer requires minimization of the output power. In the presence of another source with correlated activity, the beamformer adjusts the weight vector in such a way that, on the one hand, the gain of unity constraint is met, and on the other hand, the activity of the correlated sources is subtracted from the target activity to minimize the output power. Therefore, in the case of perfect correlation, the beamformer produces a zero SNR with respect to the activity of the target source. It should be noted that the derived rule does not depend on the topographies of the two sources.

The previous relationship was obtained under the assumption of zero additive noise 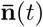. In the noisy scenario, noise covariance matrix **C**_*n*_ = *σ*^2^**I** comes into play. The sensor-space covariance matrix of observed noisy signals is given by 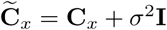. It can be shown that the estimated source variance in this case depends not only on the correlation coefficient between the two sources 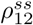 but also on the noise variance *σ*^2^ and the scalar product of source topographies 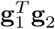:

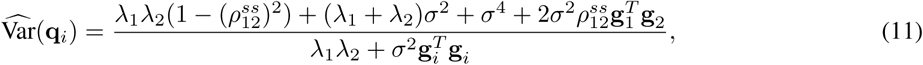

where *i* ∈ {1, 2} and *λ*_1_, *λ*_2_ are eigenvalues of matrix 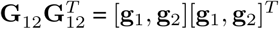.

#### 2.2.4 Example of three interacting sources

Let us now consider the environment with three active sources **q**_1_, **q**_2_, **q**_3_ with pairwise covariances 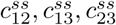. For compactness, we will assume unit variance of the source timeseries. Therefore, source-space covariance values are equivalent to the corresponding correlation coefficients 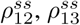 and 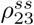. The diagram of this simulation is shown in Figure 1. Let us assume that the target source of interest is **q**_1_, but the other two sources may be also activated.

It is relatively straightforward to show, that in the case of three active unit variance sources, the LCMV estimated source power 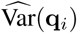 depends on the pairwise correlation coefficients as follows

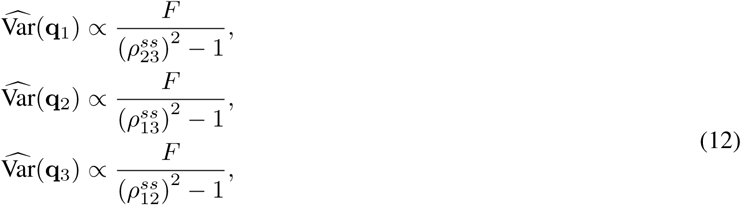

where 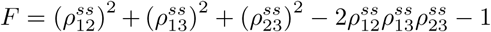.

**Figure 1:**
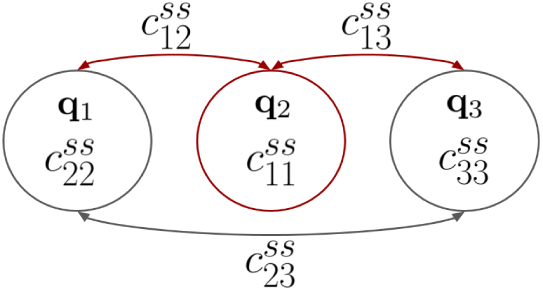
Schematic representation of three sources **q**_1_, **q**_2_, **q**_3_ power 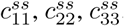. The interactions between the three sources are shown with arrows that indicate pairwise covariances 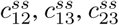. The first source **q**_1_ (highlighted in red) is the target source.

The dependence given by equation (12) can be assessed for different values of the correlation coefficients comparing 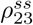 with **q**_2_ versus **q**_3_. This is shown in the heat-maps of Figure 2, where color encodes the output power of the beamformer tuned to each source (**q**_1_, **q**_2_, and **q**_3_). The power is shown as a function of the correlation between the target source **q**_1_ and the other two sources. Column of the subplots corresponds to sources and rows — to specific values of 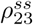. X-axis shows the correlation 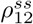 and y-axis corresponds to 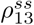. The impossible combinations of correlation values violating positive definiteness of the source-space covariance matrix are shown in white.

**Figure 2:**
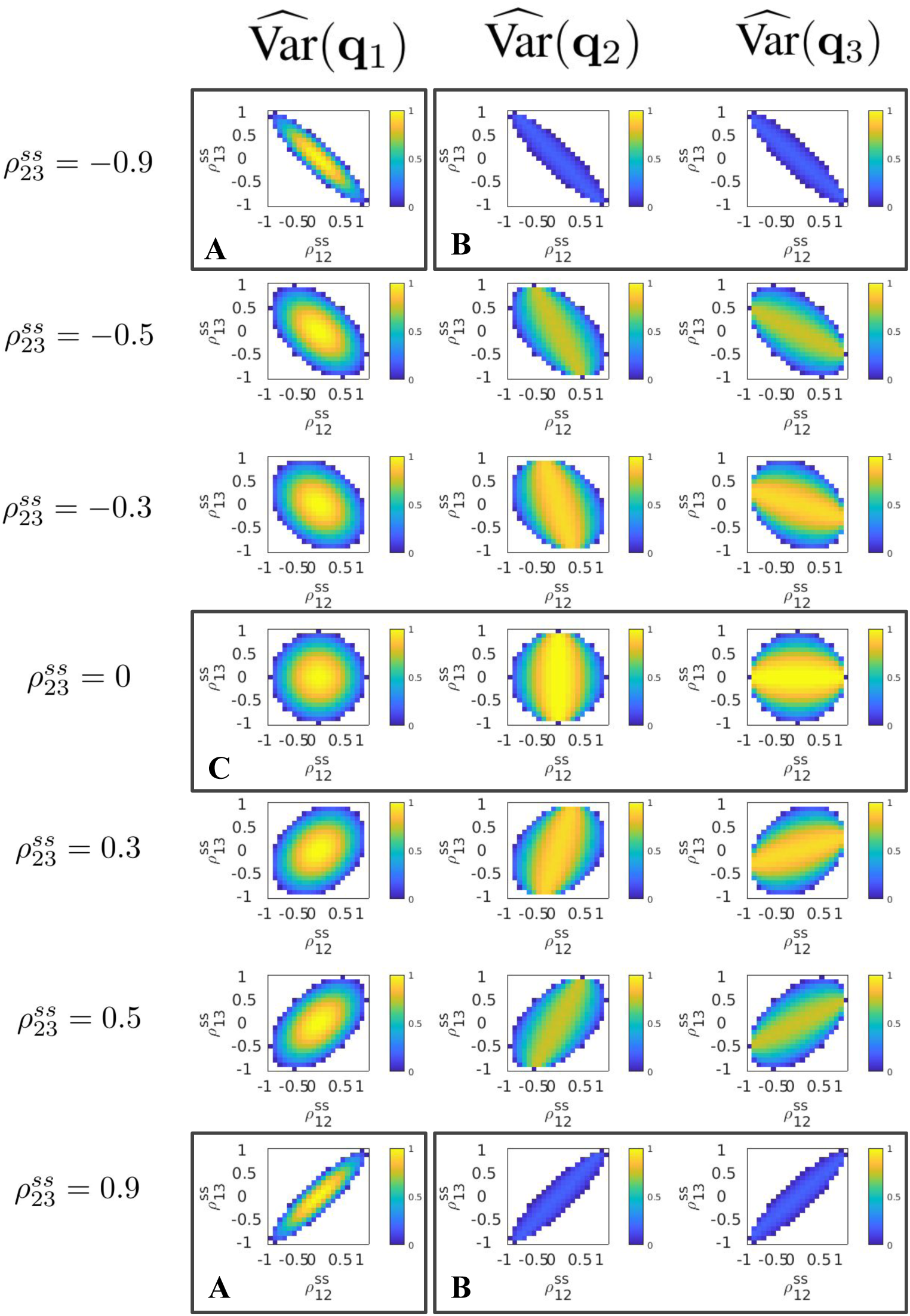
LCMV beamformer reconstruction in the case of three active sources **q**_1_, **q**_2_, **q**_3_ with pairwise correlations 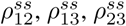 The estimated variance 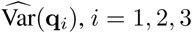 for each of three sources **q**_1_, **q**_2_, **q**_3_ as a function of pairwise correlation 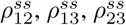 is color-coded into the heat-maps. Columns correspond to sources and row — to correlation coefficient values for the comparison between the second and third sources 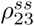. Note that the central row corresponds to the case where two of three sources are uncorrelated, 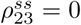. In each plot, x-axis represents correlation 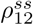 and y-axis represents 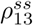. The impossible combinations of covariance values are shown in white.

It can be seen from Figure 2 that the SNR of LCMV beamformer increases with the decrease in the pairwise correlations. These combinations can be found in the middle rows of the figure, where 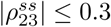, and in the central area of each subplot, where 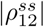 and 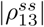 are close to zero. In the case where two sources (**q**_2_ and **q**_3_) are not coupled (Figure (2).C), the adaptive beamformer tuned to the third source (**q**_1_), the source that may become coupled to either one or both of the first two sources, remains operable for correlation coefficient up to 0.6. The variance of power estimates for the sources not coupled to any other source remains unaffected.

Secondly, a peculiar scenario is illustrated in panels Figure 2.A and B. It corresponds to the case where two (**q**_2_ and **q**_3_) out of three sources are highly correlated. This condition may occur, for example, in an auditory-motor experimental paradigm, where the subject is required to perform a motor action (e.g. press the button) in response to a deviant auditory stimulus. In this case sources **q**_2_ and **q**_3_ are located bilaterally in the primary auditory cortex and respond synchronously to the stimulus. According to our indexing, the sensory-motor response is modelled with the source **q**_1_. Under these conditions, due to the strong coupling between **q**_2_ and **q**_3_, the power of these two sources is dramatically underestimated by the adaptive beamformer. Increased correlation of **q**_1_ activity with that of the other sources further reduces the SNR of the **q**_2_ and **q**_3_ estimates. Since it is hard to detect the sources **q**_2_ and **q**_3_ under these conditions, it becomes problematic to implement the strategies suggested in (Dalal et al. (2006), Popescu et al. (2008)) to ameliorate the problems caused by source correlation.

If the activity of source **q**_1_ is completely uncoupled from that of sources **q**_2_ and **q**_3_ then, as follows from Figure 2.A, its estimate is unaffected by the extensive coupling of the other two sources. However, the range of correlation coefficients (**q**_1_ versus **q**_2(3)_) for which the estimate of **q**_1_ power remains unaffected shrinks significantly with the increase in coupling between **q**_2_ and **q**_3_. For the case of strong coupling between **q**_2_ and **q**_3_, a significant reduction in the estimate of **q**_1_ power occurs when the correlation coefficients *ρ*_12_ and *ρ*_13_ are just higher than 0.3.

Thus, we found that presence of correlated activity in the data significantly affected the estimates obtained with the adaptive beamformers, even when the source activity was only moderately correlated with the relatively strongly coupled sources.

Several solutions to the problem of correlated sources are based on setting zero constraints in the corresponding directions. (See Moiseev et al. (2011) for a review and suggestions on advancing these methods.) These solutions require localizing the jammer sources, which is a cumbersome procedure that does not guarantee success because of the likelihood of source localization errors. Additionally, these extra constraints could suboptimally consume the degrees of freedom that otherwise could be exploited for suppressing non-target activity.

In what follows we will describe a novel solution to the problem of beamforming in the presence of correlated sources. In our approach, we consider the data covariance matrix **C**_*x*_ as an element of *M* ^2^-dimensional vector space. We then employ a data independent projection procedure to mitigate the contributions from the coupled intrinsic sources.

### 2.3 ReciPSIICOS beamformer

In this section we introduce the new beamformer modification, which is immune to the contributions from correlated sources in the data. As our simulations show, our beamformer is more resistant to forward modelling errors than the classical LCMV beamformer. The key steps of the proposed algorithm are the PSIICOS projection procedure (Ossadtchi et al. (2018)) and its variations.

We originally designed PSIICOS projection technique for connectivity analysis, particularly for non-invasive detection of true zero-phase interactions between sources. The procedure employs a projection operation applied to the sensor space cross-spectral matrix treated as an element of *M* ^2^-dimensional vector space. We showed that PSIICOS could sufficiently well disentangle the subspace containing spatial leakage power from the subspace containing the contributions from the true zero-phase coupling of sources having different locations.

Here we used a similar approach to solve a complementary problem. Instead of suppressing power in the spatial leakage subspace or the subspace filled with source power, we emphasized it and instead suppressed the contributions to the correlation subspace of the sensor-space covariance matrix. We refer to the new method as ReciPSIICOS because the proposed projection is *reciprocal* to the PSIICOS projection.

#### 2.3.1 Vectorized sensor-space covariance matrix

Sensor-space covariance matrix 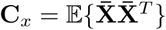 plays a pivotal role in adaptive beamforming. Here we express this matrix in terms of the underlying sources and their topographies that constitute the measured signal as defined by equation (2). By incorporating this into the MEG data generative model we obtain

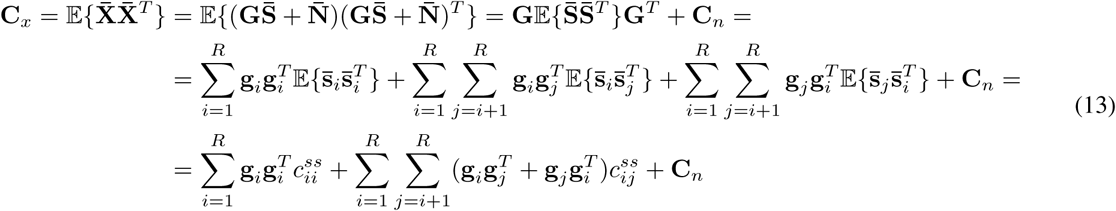

where 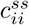 is the power of the *i*-th source, 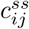 is the covariance between the *i*-th and the *j*-th source activation timeseries and **C**_*n*_ is the noise covariance matrix, which equals to identity matrix under the assumption of white noise.

We consider **C**_*x*_ as an element of *M*^2^-dimensional space and write a vectorized form of (13) to obtain

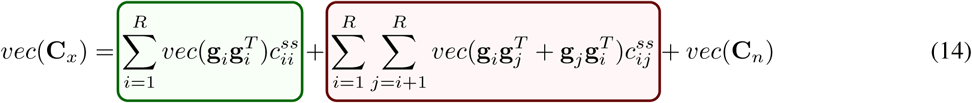

Equation (14) demonstrates that matrix **C**_*x*_ can be decomposed into two types of additive terms: auto-terms corresponding to the source power 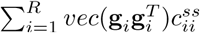 and also the pairwise cross-products of source topographies weighted with source covariance coefficients 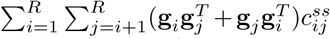. These cross-terms are present in the covariance matrix due to the non-zero off-diagonal elements *c*_*ij*_ of the source-space covariance matrix. The existence of the cross-terms in the source-space covariance matrix leads to the undesired performance of the adaptive beamformer. This happens because the weight vectors are formed in such a way that correlated sources are utilized to minimize the beamformer output power.

#### 2.3.2 Building and applying the projector

To dwindle the contribution of source covariance terms we propose implementing a projection of the vectorized data covariance matrix onto the source power subspace *𝒮*_*pwr*_ spanned by vectorized auto-products of source topographies, 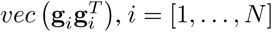, where *N* is the number of sources in the model.

It is important to notice that the subspace *𝒮*_*pwr*_ overlaps with the source correlation subspace *𝒮*_*cor*_ spanned by vectorized cross-products of source topographies 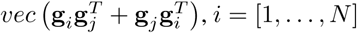 Consequently, the projection operation will inevitably suppress power in both subspaces. However, as it has been demonstrated earlier in Ossadtchi et al. (2018), the use of spectral value decomposition procedure (SVD) allows identifying a low dimensional subspace of *𝒮*_*pwr*_ capturing most of the power of the auto-product terms 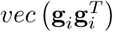. Based on this estimate of the principal power subspace, we build a projector that we then apply to the vectorized form of the observed data covariance matrix. The covariance matrix is then reshaped into the square form. Next, we proceed and design the beamformer according to the standard LCMV principles. This processing pipeline is shown in Figure 3.

**Figure 3:**
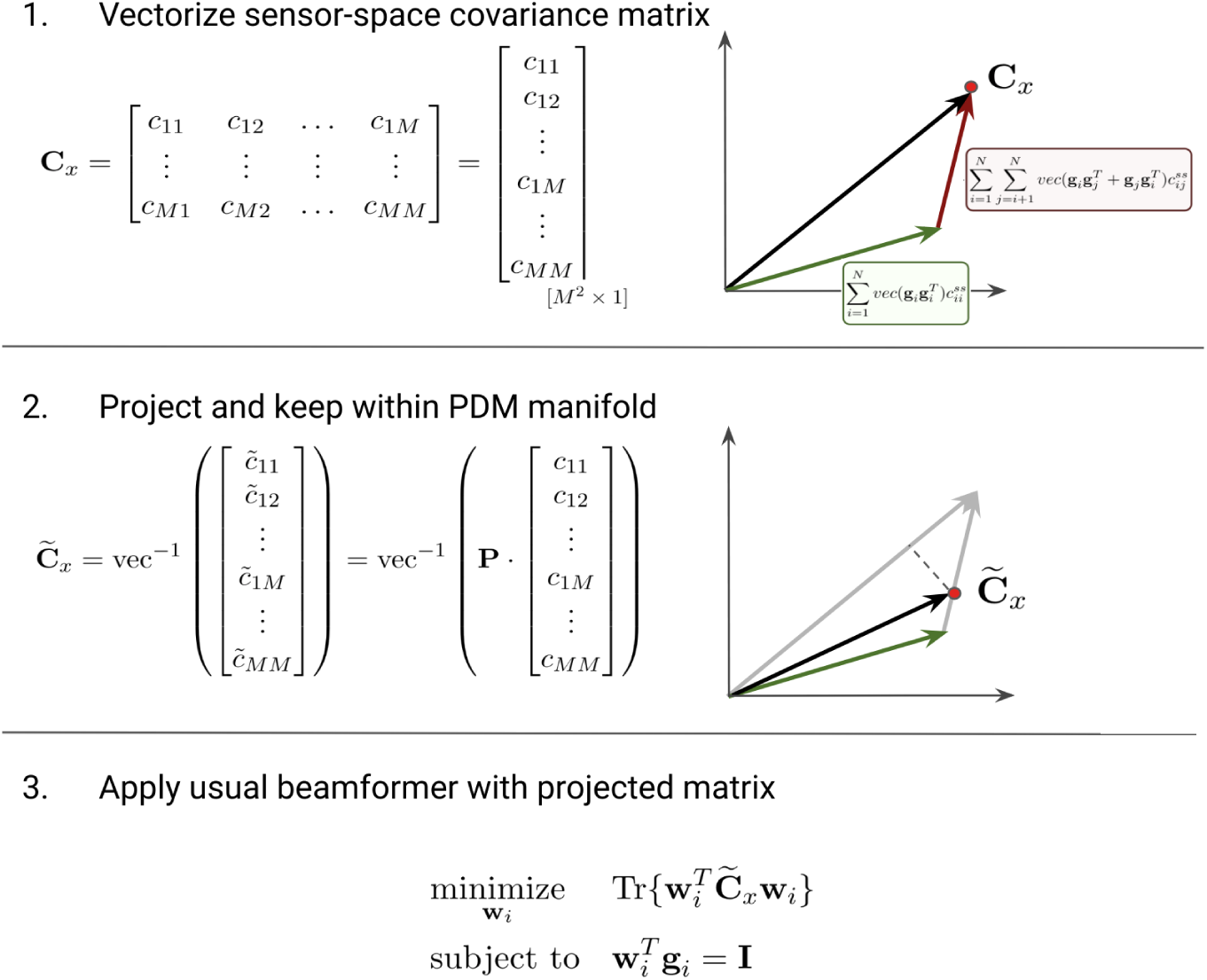
The main steps in the proposed algorithm for inverse problem solving. The sensor-space covariance matrix consists of auto-terms corresponding to the source power 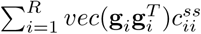 and also the pairwise cross-products of source topographies weighted with source covariance coefficients 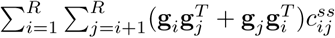. This matrix is vectorized. Next, the projection matrix **P** is calculated and applied it to the matrix **C**_*x*_ while keeping it within the positively definite matrices manifold. Finally, the usual beamformer spatial filters are calculated for the projected matrix.

To build the projector we use the following sequence of steps:

1. Construct matrix **G**_*pwr*_, where columns correspond to the vectorized auto-products of topographies for all available *N* sources from the cortical model. For compactness we are considering here the fixed orientation case and treat arbitrary source dipole orientations later in section 2.5. To abridge our expressions, we will denote the vectorized auto-product using the Kronecker product: 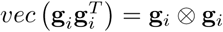. Then, matrix **G**_*pwr*_ can be created by stacking these Kronecker products as columns, i.e.

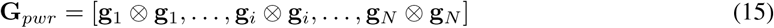
2. Compute the singular value decomposition of matrix **G**_*pwr*_ obtained on the previous step, **G**_*pwr*_ = **U**_*pwr*_**S**_*pwr*_ (**V**_*pwr*_)^*T*^, and create the projector onto the source power subspace *S*_*pwr*_

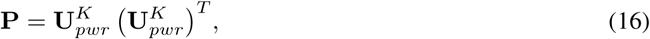

where 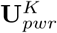 is formed from the first *K* left singular vectors. *K* is the projection rank and the only parameter which is needed to be chosen manually according to the guidelines given in section 2.4.
3. Apply the obtained projection matrix **P** to the vectorized sensor-space covariance matrix *vec* (**C**_*x*_) in order to project it onto the source power subspace and reduce the contribution of the components modulated by the off-diagonal elements of the source-space covariance matrix. The projected matrix is then

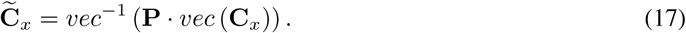
4. Since the projection procedure does not guarantee the positive definiteness of the resulting matrix 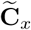, we ensure this property by replacing its negative eigenvalues with their absolute values. In other words, we calculate the final data covariance matrix with suppressed contribution of the correlated sources as

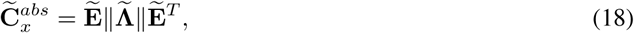

where 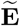 and 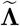 are the matrices containing eigenvectors and eigenvalues of the matrix after the projection 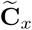.
5. Use projected covariance matrix 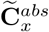 in (4) instead of the original **C**_*x*_ to compute spatial filters **w**_*i*_, in (5) to reconstruct the source activation timeseries and in (6) to estimate the source power 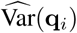 distribution.

The ReciPSIICOS procedure of source reconstruction is schematically shown in Figure 4 for the case of ERP analysis. We use MEG data projected into the forward model principal subspace defined as the subspace capturing 90% of the energy in the forward operator matrix. This allows reducing the number of dimensions to about 60 to 80 virtual sensors. Following the preprocessing, we split the data into the epochs according to the timestamps of the stimulus onsets. Next, we calculate average evoked response (ERP). We then use these average ERPs to compute sensor-space covariance matrix. At this step, we have to make sure that the ERP curve is of sufficient length and that our data covariance matrix is full rank.

**Figure 4:**
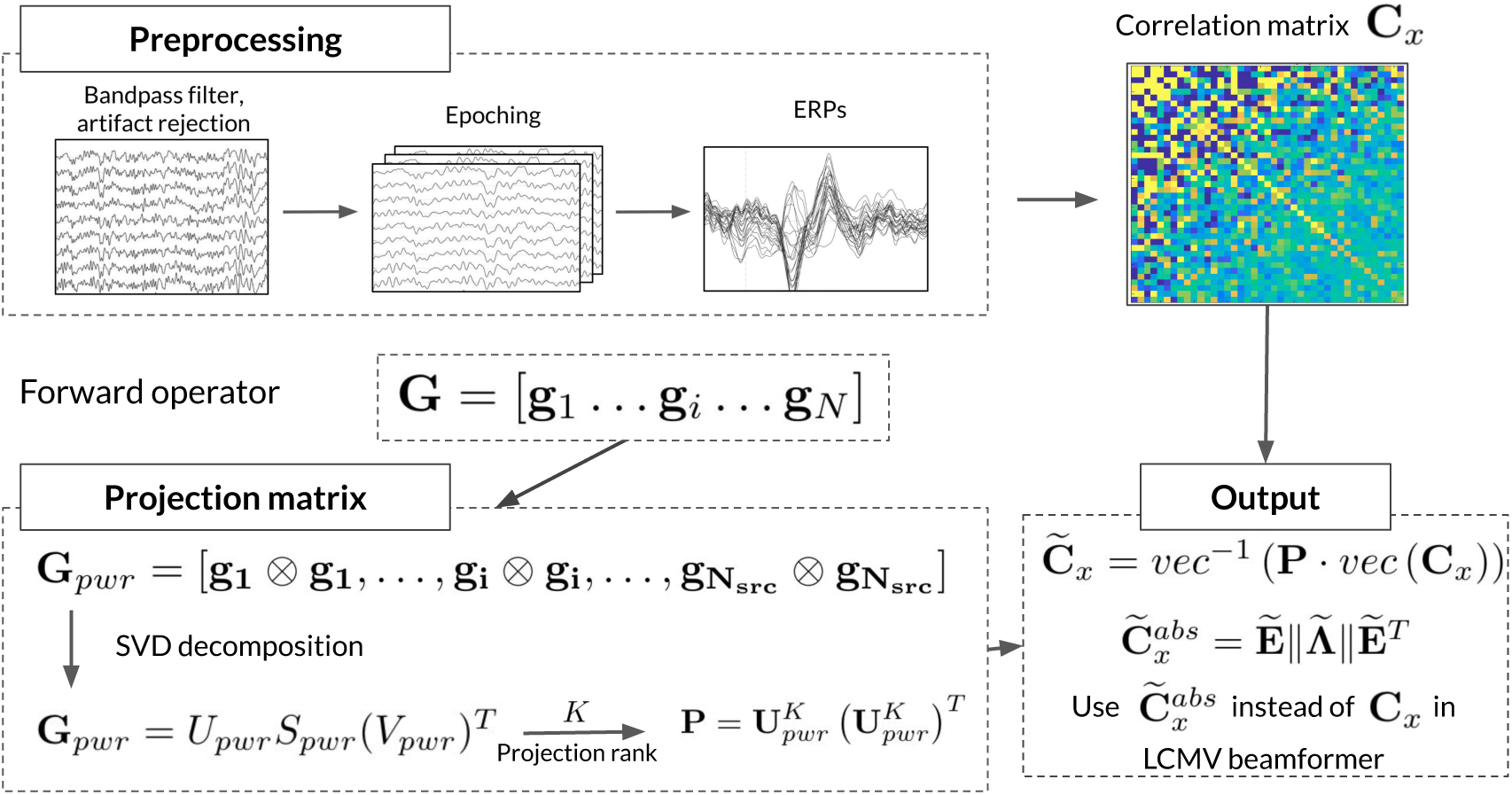
Schematic presentation of the ReciPSIICOS algorithm.

We follow the steps listed in the beginning of this section to prepare projection operator based on the forward model matrix corresponding to the virtual sensors. This has to be done only once for a given forward model matrix. If the forward model matrix needs to be recomputed for some reason, the ReciPSIICOS projector has to be recalculated.

We then apply this projection operator to the vectorized data covariance matrix and reshape the result of the projection to a square matrix and heuristically fix negative eigenvalue problem as described above. Our experience shows that contribution of the eigendirections modulated by the negative eigenvalues is quite slim and does not exceed 20% of the total energy in the eigenvalue spectrum of the projected matrix (see Figure 21). We therefore believe that the proposed heuristics provides an adequate balance of efficiency and simplicity as compared to the full blown consideration of Riemannian geometry of the correlation matrices, which nevertheless constitutes an interesting direction for further improvement of our method.

In this section we have described a simple projection into the power subspace. This method delivers a reasonable performance and provides adaptive beamforming scheme with immunity to the source-space correlations inevitably present in the MEG and EEG data. See section 3 for description of simulations and real-data analysis results where we refer to this method as ReciPSIICOS.

Because *𝒮*_*pwr*_ and *𝒮*_*cor*_ subspaces overlap, the described projection procedure depletes power from both subspaces, but does so at a faster pace for *𝒮*_*cor*_ subspace. The ultimate balance between the power in the two subspaces achieved with specific value of projection rank *K* may serve as a quality metric for the procedure. We use these considerations in section 2.4 where we suggest a strategy for choosing optimal projection rank.

In what follows we will describe another, slightly more complicated procedure, that allows achieving a better balance between the depletion ratio of power in the *𝒮*_*pwr*_ and *𝒮*_*cor*_ subspaces.

#### 2.3.3 Projection away from correlation subspace whitened with respect to the power subspace

The idea behind this procedure lies in developing a projector not into the *𝒮*_*pwr*_ subspace, but rather away from the *𝒮*_*cor*_ subspace. In addition, we perform this projection in the space whitened with respect to the power subspace, which ensures that minimal possible power will be depleted from *𝒮*_*pwr*_. In what follows we refer to this method as Whitened ReciPSIICOS (WReciPSIICOS). Step-by-step algorithm of building and applying the whitened projector is presented below as well as the schematic representation in Figure 5.

**Figure 5:**
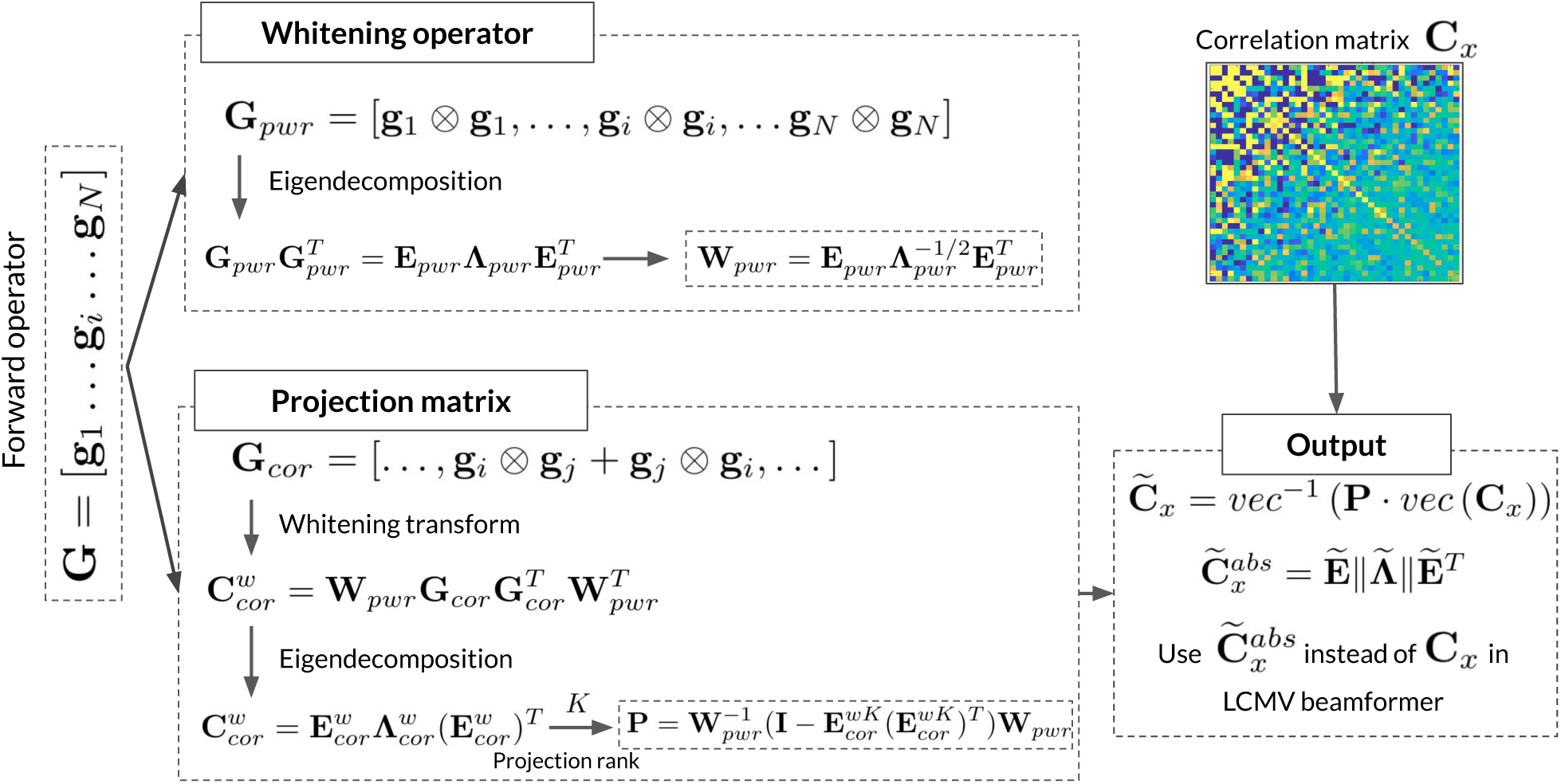
Schematic presentation of the Whitened ReciPSIICOS algorithm.

1. Construct matrix **G**_*cor*_ whose columns span correlation subspace *𝒮*_*cor*_. The columns of this matrix contain vectorized sums of symmetric source topographies outer-products for all 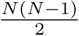 ordered pairs of sources

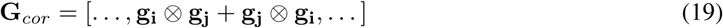

and calculate matrix 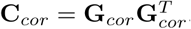
2. Construct matrix **G**_*pwr*_ whose columns span power subspace *𝒮*_*pwr*_. Columns of this matrix are the vectorized sums of source topography vectors as in the previous projector

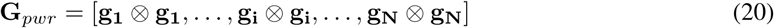

Then calculate the matrix 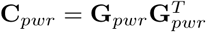.
3. Using eigendecomposition of **C**_*pwr*_, compute the whitening operator **W**_*pwr*_ for the source power subspace *𝒮*_*pwr*_

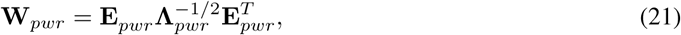

where **E**_*pwr*_ is the matrix of eigenvectors of **C**_*pwr*_ and diagonal matrix **Λ**_*pwr*_ contains the corresponding eigenvalues.
4. Apply whitening transformation to **C**_*cor*_ to obtain 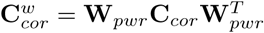.
5. Extract the principal subspace of 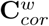 by means of eigenvalue decomposition as

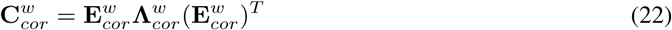
6. Form matrix projecting away from the source correlation subspace *𝒮*_*cor*_ and operating in the space whitened with respect to *𝒮*_*pwr*_

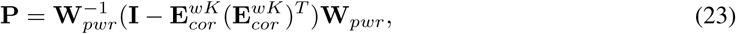

where **I** is identity matrix, 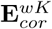 is the matrix of the first *K* eigenvectors of matrix 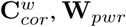 is the whitening matrix computed earlier and *K* is the projection rank and the only parameter in the introduced algorithm that needs to be chosen manually.
7. Apply the obtained matrix **P** to the vectorized sensor-space covariance matrix *vec* (**C**_*x*_) in order to project it onto the source power subspace and reduce the contribution of the components modulated by the off-diagonal elements of the source-space covariance matrix. The projected matrix is then

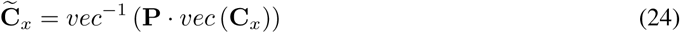
8. Since this whitened projection procedure also does not guarantee the positive definiteness of the resulting matrix 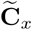, we ensure this by replacing its negative eigenvalues with their absolute values. In other words, we calculate the final data covariance matrix that has the contribution of the correlated sources suppressed as

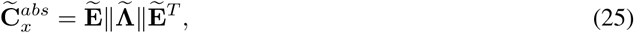

where 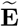 and 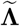 are the matrices containing eigenvectors and eigenvalues of 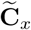.
9. Use the projected covariance matrix 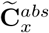 in (4) instead of the original **C**_*x*_ to compute the spatial filters **w**_*i*_ to reconstruct the source activation timeseries ((5)) and to estimate the source power 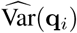 distribution (in (6)).

As in the previous case, this projection procedure is designed without taking into account the Riemannian geometry of the manifold of the correlation matrices. Interestingly, this approach is based on projecting the original vectorized covariance matrix away from *𝒮*_*cor*_ yields even smaller fraction of negative eigenvalues. In our experience, this whitened projection operator consistently results into less the 20% of the total energy in the eigenvalue spectrum of the projected matrix (see Figure 21).

### 2.4 Optimal projection rank

In the previous sections we introduced two projection procedures aimed at suppressing the contribution of the components modulated by the off-diagonal elements of the source-space covariance matrix. The projection procedures require only one parameter, the projection rank, that has to be preset by the operator. Since the projection depends only on the forward model, a possible solution is to select the projection rank *K* based on the simulations like the ones described in section 3.1.

The other approach is based on the following considerations. We operate in the *M* ^2^-dimensional space and consider equation (14) as the observation equation of the *vec*(**C**_*x*_). This equation represents *vec*(**C**_*x*_) as a superposition of activity of sources with the topographies 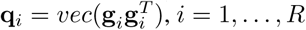, *i* = 1, …, *R* and 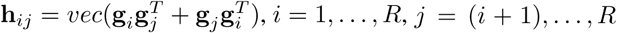, *i* = 1, …, *R, j* = (*i* + 1), …, *R*. Vectors **q**_*i*_ and **h**_*ij*_ span *𝒮*_*pwr*_ and *𝒮*_*cor*_ subspaces correspondingly. To calculate maximum possible energy in each of the subspaces we can assume that the “sources” with “topographies” **q**_*i*_ and **h**_*ij*_ are activated with unit independent activations and then the total energy in both subspaces can be measured as the trace of the *M* ^2^ × *M* ^2^ correlation matrices **C**_*pwr*_ and **C**_*cor*_ introduced earlier in section 2.3.3. After the projection procedure of rank *k* is performed by means of matrix **P**_*k*_, the subspace correlation matrices of the observable 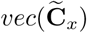 will read as 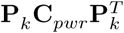 and 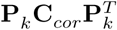. The fraction of power left in both subspaces can be then computed as

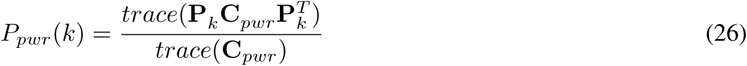

and

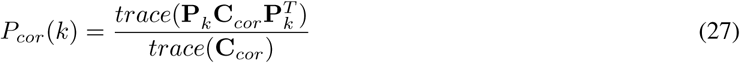

Based on these equations we can plot the curves (*P*_*pwr*_(*k*), *P*_*cor*_(*k*)) parameterized by projection rank parameter *k*. An example of such a plot is shown in Figure 6 that depicts the fraction of past projection power in both subspaces parameterized by the projection rank *k*.

**Figure 6:**
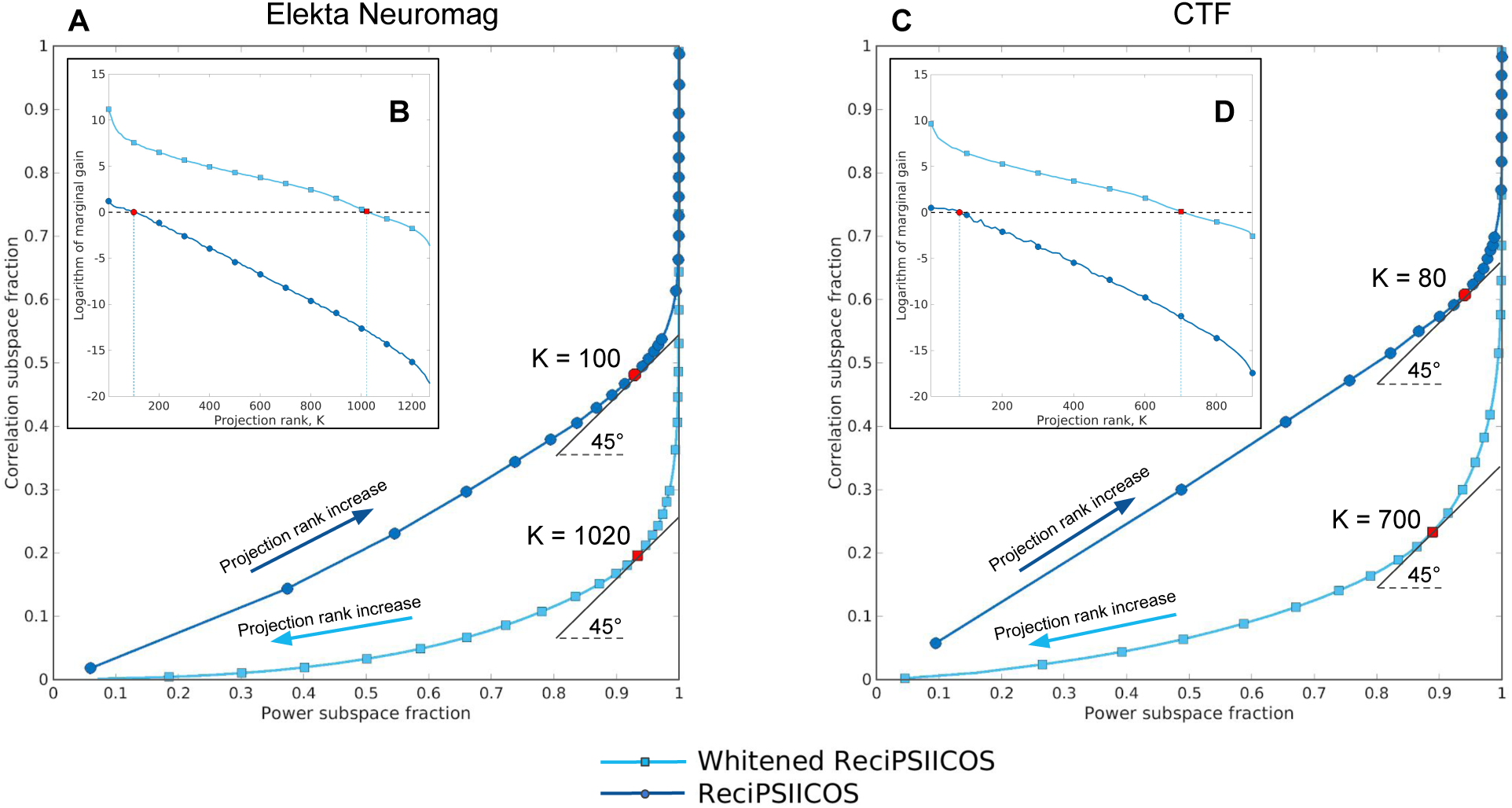
Parametric curves of power depletion in the *𝒮*_*pwr*_ and *𝒮*_*cor*_ subspaces parameterized by projection rank for ReciPSIICOS and Whitened ReciPSIICOS projections for Elekta Neuromag (panel A) and CTF (panel C) machines. Panels B and D show the logarithm of marginal gain, 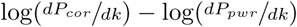, as a function of projection rank. The optimal projection rank is the value which sets the logarithm of marginal gain in power depletion to zero, which corresponds to the 45°angle of power suppression curves tangent at this point.

With the reduction of projection rank in the ReciPSIICOS procedure that performs projection onto the power subspace, more power is depleted from both the subspaces *𝒮*_*pwr*_ and *𝒮*_*cor*_. Conversely, the Whitened ReciPSIICOS implements the projection away from the correlation subspace, so the power is suppressed while the projection rank grows.

Our method works because there exists a range of projection rank values, where the power is depleted faster from the correlation subspace than from the power subspace, i.e. 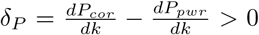. However, when projection rank *K\** reaches a certain value, the rate of depletion of power from the two subspaces become comparable. Consequently, for the values of *k* > *K\**, the difference *δ*_*P*_ changes its sign to negative, i.e. the power subspace starts to lose power faster than the correlation subspace. This projection rank value *K\** can be considered as optimal. Our simulations show that the projection operation remains fairly stable for a broad range of projection rank values *k*. Yet, these values are affected by the number of virtual sensors used and would significantly differ for different MEG probes.

Figure 6 demonstrates the power suppression curves for the two proposed methods and for the two MEG arrays: 204-gradiometer array of Neuromag Vectorview-306 (panel A) and 275 gradiometer array of CTF (panel C). In both cases, the number of virtual sensors was selected so that their leadfiled matrix captured 99% of variance present in the original leadfield matrix which resulted into *M*^*Nmg*^ = 50 and *M*^*CTF*^ = 42 number of sensors. Panels B and D show the logarithm of marginal gain, defined as 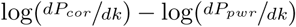, as a function of projection rank. According to the imposed assumptions, the optimal projection rank is the value which sets the logarithm of marginal gain in power depletion to zero, which also means that the angle of the tangent to power suppression curve is equals to 45°at this point.

This approach demonstrates the clear superiority of Whitened ReciPSIICOS method over plain ReciPSIICOS, as for each value of preserved power in *𝒮*_*pwr*_ Whitened ReciPSIICOS suppresses more power in *𝒮*_*cor*_.

### 2.5 Handling unknown source orientations

Anatomically, the dipole orientations coincide with the direction of apical dendrites of the pyramidal neurons and therefore are predominantly orthogonal to the cortical mantle. Modern tools of MRI data analysis allow for a very accurate extraction and precise parametrization of the cortical surface with the number of nodes on the order of tens of thousands, which in turn results in a reasonable accuracy of the orientation specification. The uncertainty that remains may be efficiently modeled with such techniques as described in Lin et al. (2006).

Because of the memory and processing time limitations when performing exploratory source-space synchrony analysis, we have to use a significantly downsampled version of the cortical mantle that contains only several thousands of nodes. Such downsampling drastically reduces the requirements to computational resources but introduces a significant uncertainty in the orientation of elementary sources.

It has become a common practice to restrict the source space to the downsampled cortical mantle and to model the arbitrary orientations of node dipoles by representing each elementary source with the triplet of orthogonal dipoles. In case of MEG, the magnetic field is equal to zero outside the spherical conductor produced by a dipole with radial orientation. Because of this, the triplet can be efficiently replaced by a pair of dipoles in the tangential plane calculated for each node.

For an arbitrary orientation vector at some *i*-th vertex 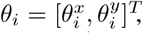 the corresponding dipole topography is 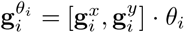, where 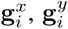 are the topographies of the two orthogonally oriented dipoles in the tangential plane at the *i*-th vertex. By varying the orientation angle we can obtain an infinite set of power subspace topography vectors 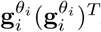.

By expanding these auto-products of topographies of the oriented dipoles in terms of the topographies 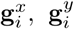, *i* = [1, …, *N*] oriented along *x* and *y* axis of the local tangential plane we obtain

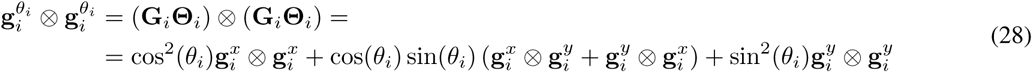

Therefore, in order to accommodate the arbitrary orientation constraint, elements **g**_*i*_ ⊗ **g**_*i*_ in equations (15) and (20) have to be replaced by 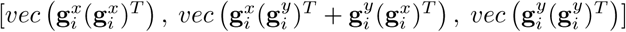.

Arbitrary orientations lead to a slightly more cumbersome manipulation for the cross-products of topographies. Similarly to the auto-products step, sums of Kronecker cross-products of topographies of different arbitrarily oriented sources present in equation (19) can be expanded as

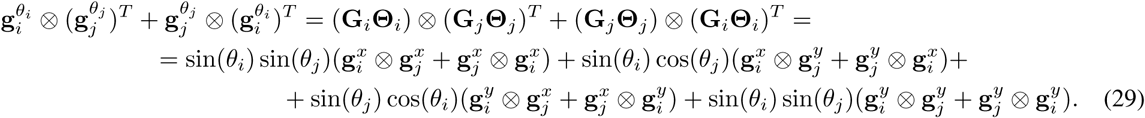

Therefore, elements (**g**_*i*_ ⊗ **g**_*j*_ + **g**_*j*_ ⊗ **g**_*i*_) in equation (19) are to be replaced with

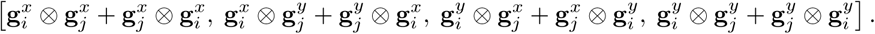

The rest of the procedure for building the projector is unaltered.

### 2.6 Monte Carlo simulations

In order to compare the proposed algorithms with other relevant source reconstruction methods we ran several Monte Carlo simulations. To simulate the MEG signals, we used cortical surface model with 20000 vertices reconstructed from anatomical MRI data using FreeSurfer software (Fischl (2012)). The forward model matrix **G** for freely oriented dipolar sources was computed with Brainstorm software (Tadel et al. (2011)) using the overlapping spheres procedure. Each location on the cortical grid was served by two topography vectors confined to the locally tangential plane as determined by the first two right singular vectors of the local [*M* × 3] forward matrix.

In different experiments, we studied the interaction of two or three sources. For each Monte Carlo trial, a random set of pairs or triplets of dipolar sources was picked as the target stimulus-related sources. One hundred epochs were generated and then averaged to obtain the event related field (ERF). The activations of target sources **s**(*t*) were modeled with 10-Hz sinusoids.

The simulated data mimicked two experimental conditions: the first with highly correlated source activation timeseries and the second with uncorrelated ones. To model these two extreme cases, the phase difference between activation timeseries was set to zero for the correlated condition and to 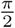 for the uncorrelated condition. Each trial onset was jittered with respect to the task onset by adding a random shift generated from zero-mean normal distribution with the standard deviation corresponding to 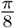 phase difference.

We modeled task irrelevant activity with *Q* = 1000 spatially coherent, task-unrelated cerebral sources whose locations and timeseries varied with each epoch. Source locations were mapped on the nodes of the high resolution cortical grid (20000 vertices). The activation timeseries were narrow-band signals obtained via zero-phase filtering of the realizations of Gaussian (pseudo)random process by the fifth order band-pass IIR filters in the bands corresponding to theta (4-7 Hz), alpha (8-12 Hz), beta (15-30 Hz) and gamma (30-50 Hz, 50-70 Hz) activity. Their relative contributions were scaled in accordance with the well-known 1*/f* characteristic of the MEG spectrum. We scaled the brain noise components to match typical signal-to-noise ratio of real-life recordings. To project these sources into the sensor space, the corresponding columns of the forward matrix was computed for the high resolution source grid. We simulated 100 epochs of ERF data and for each epoch a new randomly picked set of noisy sources was chosen and new noisy timeseries with approximately 1*/f* spectrum were generated. We defined the SNR in our simulated data in the sensor space as the ratio of Frobenius norms of data matrices for the induced and brain noise components filtered in the band of interest (0.5-7 Hz), corresponding to the ERF response.

The high resolution grid of 20000 vertices was used only for data simulation. For the source reconstruction process we employed a 4 times sparer cortex grid with 5000 vertices. We ran 500 simulations and compared two versions of the proposed ReciPSIICOS method against the classical adaptive LCMV beamformer and MNE techniques.

### 2.7 Performance metrics

In our Monte Carlo simulation analysis, goodness of localization was estimated using three metrics: source localization bias, point spreading radius and ratio of successful detection. While the data was simulated using dense cortical model with 20000 sources, source reconstruction procedure was based on the sparse cortical model with 5000 sources to make the reconstruction procedure more realistic. We considered separately the scenarios with two and three target sources. In order to run the comparative analysis for each Monte Carlo trial we repeated the following steps:

1. Given the simulated data with target sources 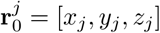, where *j* = {1, …, *N*_*src*_} for number of sources *N*_*src*_ = 2 or *N*_*src*_ = 3, estimate the power of each source **Z** = [*z*^1^, …, *z*^*M*^] using the four studied methods (ReciPSIICOS, Whitened ReciPSIICOS, LCMV, MNE). The sources with estimated power below the threshold *α* are considered as inactive. *α* is computed as a fraction of maximal estimated power value among all sources as *α* = *a* max(**Z**), 0 ≤ *a* ≤ 1. To mimic real-life situation, we choose the parameter *a* individually for each Monte Carlo iteration and each method, according to the following procedure. We scan through the grid of different values of *a*. Each threshold value results into a specific number of connected regions (clusters). We than choose the highest value of the threshold that results into the number of sources that were simulated *N*_*src*_. In case we can not find such a threshold value, we repeat the above procedure for *N*_*src*_− 1 sources. Sources with estimated power exceeding *α* are considered as active and assigned to the closest target source in terms of the Euclidean distance.
2. All active sources are divided into *C* clusters 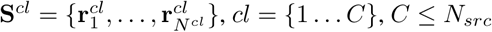, according to their correspondence to one of the target sources, where *N*^*cl*^ is the number of sources in each cluster. In each cluster find the source with maximal estimated power

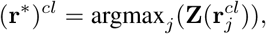

where *j* ∈ 1 …*N*^*cl*^, *cl* = {1, 2} or *cl* = {1, 2, 3}.
3. Calculate the average Euclidean distance between the local maximums in each cluster and the corresponding target source. The obtained value reflects the source localization bias and is measured in meters:

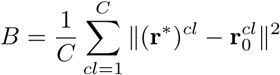
4. To estimate the point spreading radius, first normalize the estimated power values inside each cluster

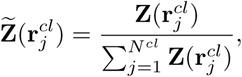

where *j* ∈ 1 …*N*^*cl*^, *cl* = {1 …*C*}, and then calculate the distance between each source belonging to the current cluster and the source with maximal estimated power. The point spreading radius is then the averaged sum of computed distances weighted with the normalized power values

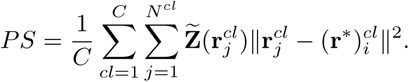
5. If the number of detected clusters *C* is lower than the number of modeled sources (2 or 3), this simulation is considered as an unsuccessful detection. Also, if the source localization bias is higher that 2cm, which means that the peak of reconstructed activity is 2 cm away from the initial location, this case is also considered as an unsuccessful detection. The detection ratio is then defined as the ratio of successful detection across all simulations.

For all simulations, we computed statistical distributions of the metrics of interest (9) and the corresponding median values (8). We used localization bias and point spreading area as accuracy metrics for each method.

The described procedure and considered metrics give the highly intuitive output and result into criteria which are usually used in the manual expert evaluation of source reconstruction results, that is why we use them for comparison. Moreover, the obtained characteristics of LCMV and MNE reconstruction perfectly fit to the previous results (Rana et al. (2018)), which also confirms the consistency of the chosen metrics.

### 2.8 MEG data acquisition and handling

We applied the ReciPSIICOS and Whitened ReciPSIICOS algorithms to the MEG datasets from two experiments that involved auditory processing. The behavioral tasks and MEG recording systems were different in these experiments.

#### 2.8.1 Dataset 1

The first dataset contained MEG data for one healthy subject and one ASD patient. The data were collected at Moscow MEG facility with Elekta-Neuromag Vectorview 306 system (Elekta Oy, Finland) with 204 planar gradiometers and 102 magnetometers. The subjects participated in passive listening session. During the experiment, the 40 Hz auditory stimuli sequence was presented monaurally in the left ear. The session consisted of 80 trials. MEG data was recorded with 1000 Hz sampling frequency. The data was preprocessed using the Elekta MaxFilter™ software. To compute the auditory steady-state responses (ASSR), we considered the low gamma-band oscillations as the primary focus of the analysis. The preprocessed data was band-pass filtered in the 35-45 Hz band using a band-pass FIR filter with 128 taps. The obtained signals were epoched 500 ms prior- and 1s post-stimulus and averaged to obtain the ERF.

The sensor-space covariance matrix was computed for the average ERF. The length of the ERF timeseries (including the pre-stimulus interval) was 1.5 s (1500 samples) for the virtual sources count of *M* = 50. This number of samples was sufficient for the estimation of the sensor-space covariance matrix. Based on the individual 1.5 T MRI scans forward model, matrices comprising topographies of freely oriented dipoles in the 20000 mesh grid nodes were calculated individually for the two participants.

#### 2.8.2 Dataset 2

The second dataset was collected in one subject using a CTF-275 MEG system. The data was recorded in the Montreal Neurological Institute, McGill University, Canada by Elizabeth Bock, Peter Donhauser, Francois Tadel and Sylvain Baillet for Brainstorm tutorial (Tadel et al. (2011)). The subject received auditory stimuli binaurally through intra-aural earphones (air tubes+transducers). The stimuli included a total of 200 regular beeps (440 Hz) and 40 easy deviant beeps (554.4 Hz, 4 semitones higher). The inter-stimulus interval was randomized and uniformly sampled between 0.7 s and 1.7 s seconds. The subject was in a seating position. He was instructed to press a button with the right index finger when detecting a deviant stimulus. Auditory stimuli were generated with the MATLAB Psychophysics toolbox. MEG data was acquired at 2400 Hz. The anti-aliasing low-pass filter with cutoff frequency of at 600 Hz was applied and the data was saved with the 3rd order gradient. Eye artifacts were removed using ICA decomposition. The data was bandpass filtered in the 1-70 Hz band using a band-pass FIR filter with 128 taps. The obtained signals were epoched 100 ms prior- and 500 ms post-stimulus and averaged separately for deviant and standard stimuli. To obtain an MMNm component, we computed the difference between the deviant and standard responses.

#### 2.8.3 Forward and inverse operators

Forward operators were computed via Brainstorm software (Tadel et al. (2011)) using an overlapping spheres model. MNE and LCMV inverse operators were computed in agreement with their implementation in the Brainstorm software. Due to the typically high SNR in the ERF/ERP data, no noise covariance was supplied to either of the algorithms. ReciPSIICOS and Whitened ReciPSIICOS inverse operators, as well as all the simulations and analysis, were performed using custom MATLAB scripts.

## 3 Results

### 3.1 Monte Carlo simulation study

The performance of ReciPSIICOS and Whitened ReciPSIICOS projections was tested using Monte Carlo simulations, according to the methodology described in section 2.6. To perform a comparative analysis for each simulation, the source power was estimated using four techniques discussed above: ReciPSIICOS, Whitened ReciPSIICOS, LCMV and MNE. Simulated data were generated using a dense cortical grid of 20000 sources, and source reconstruction procedure used a four times sparser cortical model with 5000 sources. This study design allowed to perform a more realistic simulation analysis, though naturally entails small additional persistent bias in all considered metrics.

#### 3.1.1 Two interacting sources

Source reconstruction results for a single trial of Monte Carlo simulations with two active sources are shown in Figure 7. The left and right panels show the results for the synchronous and asynchronous pairs of sources, correspondingly. For each Monte-Carlo iteration, such as the one depicted in Figure 7, the pair of symmetrically located dipolar sources was picked randomly. SNR in the ERP corresponding to the simulated data was set to 4. Each subplot on the graph demonstrates cortical distribution of the estimated variance and the scatter plots allow to appreciate the dynamic range of the obtained solutions. As expected, in the case of the asyncronous sources, ReciPSIICOS, Whitened ReciPSIICOS and LCMV demonstrated spatial super-resolution and very tight activation distributions, while MNE-produced map was characterized by a significantly greater cortical spread. For synchronous sources, the LCMV beamformer clearly showed a signal cancellation effect and completely failed to localize true sources, while the proposed projection remained operable in both ReciPSIICOS and Whitened ReciPSIICOS versions.

**Figure 7:**
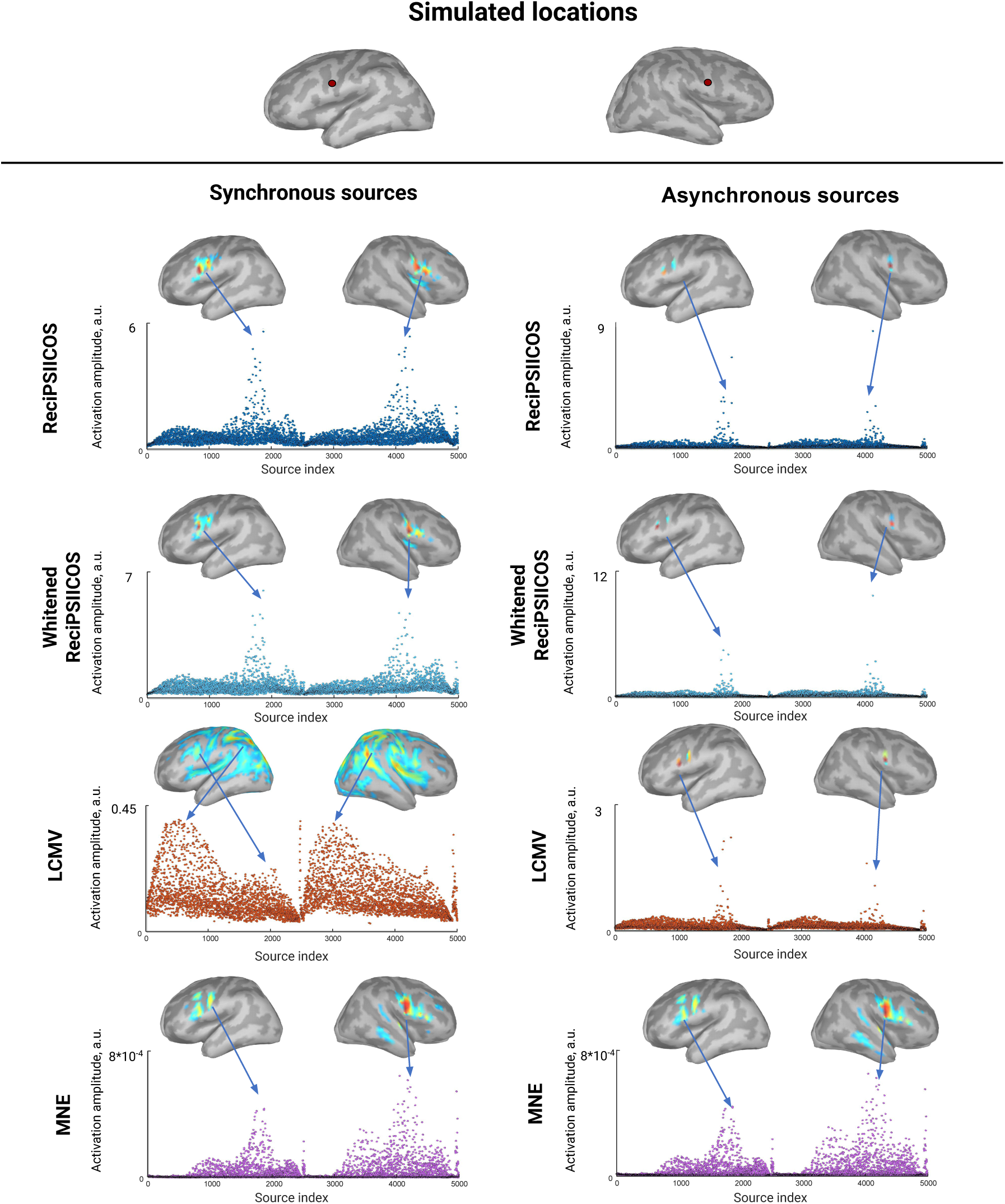
Estimated source power for one Monte Carlo simulation. The pair of active symmetric sources was picked randomly and activated synchronously (left panel) and asynchronously (right panel). The source reconstruction was performed using four techniques: ReciPSIICOS, Whitened ReciPSIICOS, LCMV and MNE. The noise was set to SNR = 4. Each subplot shows the estimated amplitudes (y-axis, scale is individual) for each of 5000 sources (x-axis).

It is important to note that, as evident from this single MC-trial example (Figure 7), the proposed projection did not adversely affect the super-resolution property of the beamformer-based inverse solution in the asynchronous case and kept operating in the scenario with synchronous sources whereas the original adaptive LCMV failed to recover the simulated sources. The MNE technique was insensitive to the correlation between sources and showed qualitatively identical results in both synchronous and asynchronous cases. We would also like to point out that the the magnitude of and dynamic range of ReciPSIICOS and Whitened ReciPSIICOS obtained solutions appeared to be noticeably greater than those delivered by the MNE and the LCMV techniques.

As we will show in the later sections, the norms of the inverse operator rows for the LCMV and the two techniques proposed here are of comparable magnitude. Therefore the increased magnitude of the output in both synchronous and asynchronous cases is explained by the proper orientation of the corresponding spatial filter (row of the inverse operator) aimed at emphasizing target source activity and not using activation of synchronous sources to chase the minimum variance requirement. The fact that the increase in magnitude was observed in both synchronous and asynchronous cases can be explained by the fact that the sources present in the realistically simulated brain noise are used by the greedy LCMV solution to further minimize the output variance.

To generalize the observations described above we performed 500 Monte Carlo simulations with 500 trials and used the three metrics detailed in section 2.7: source localization bias, radius of point spreading and ratio of successful detection to compare the performance of the four methods. We investigated the performance for various SNR and forward model inaccuracies and presented them as medians computed over all 500 MC iterations for each condition.

Figure 8 demonstrates the results of comparative analysis of performance as a function of SNR (panel A) and forward model inaccuracies (panel B). Consider first the noise-free forward model operator (panel A). All four methods, except for the adaptive LCMV in the synchronous case, demonstrate stable performance for SNR > 2. In case of uncorrelated source activations, ReciPSIICOS and Whitened ReciPSIICOS, as well as LCMV beamformer, demonstrate high performance: detection ratio is close to 100%, localized area is compact and coincides with the simulated location. At the same time, MNE demonstrates low localization bias and, as expected, a greater point spreading radius. It detects only one source for 40% of trials MNE. These localization characteristics perfectly fit the findings described in the recent study Rana et al. (2018).

**Figure 8:**
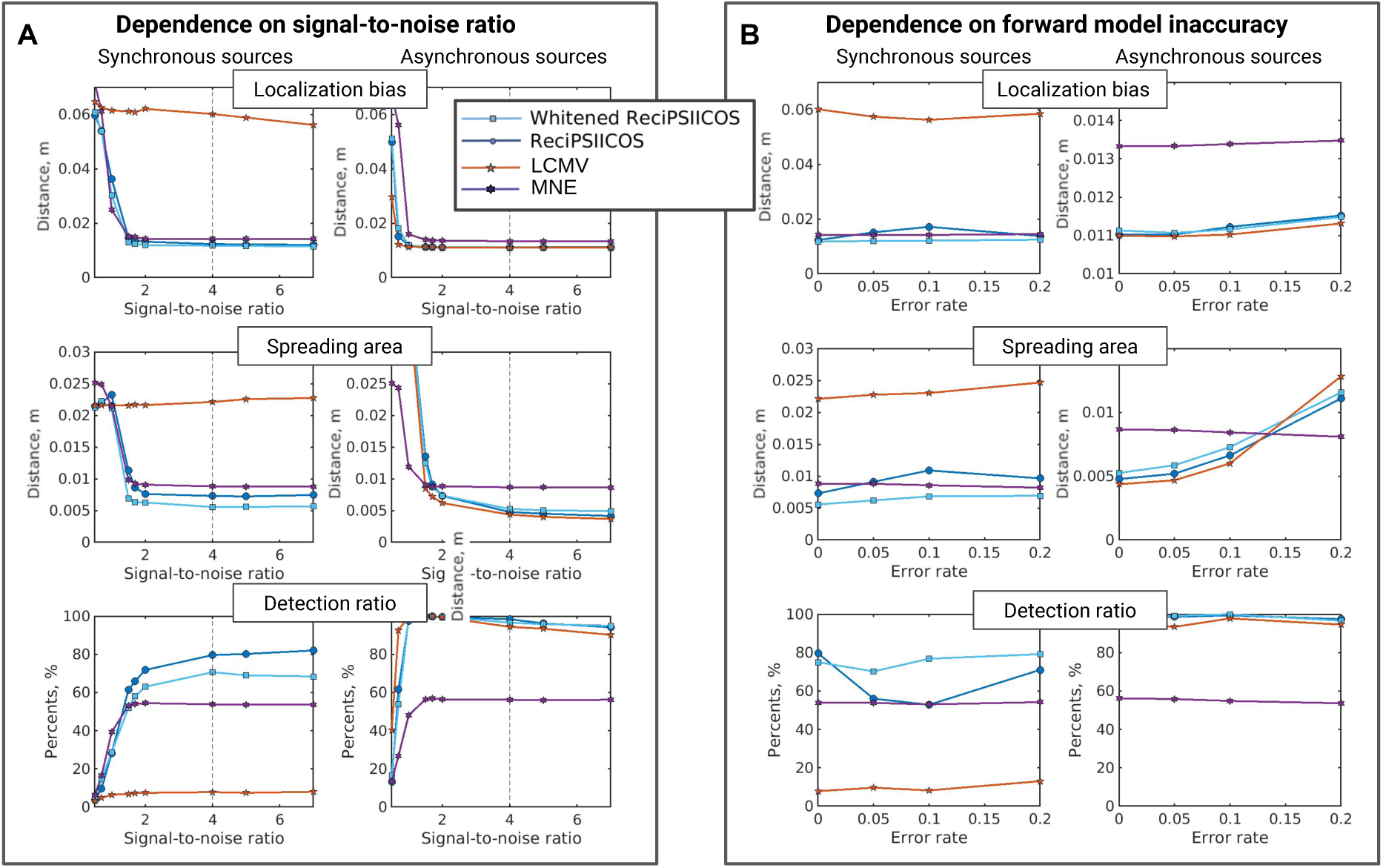
Monte Carlo simulation study results. Comparison of ReciPSIICOS, Whitened ReciPSIICOS, LCMV and MNE localization for synchronous and asynchronous sources according to three metrics: localization bias, area of point spreading and detection ratio. **A**. Dependence of estimated metrics on SNR. **B**. Dependence of estimated metrics on forward model inaccuracy.

Considering the right panel of Figure 8.A, one can see that for correlated signals LCMV performance drops significantly and does not improve with the growing SNR. Due to severe signal cancellation the detection ratio characteristics is below 10%, which means that only in 10% of cases LCMV detects two activation blobs with the maximum value not further than 2 cm from the simulated locations. While the three methods reach equally good localization bias at the level comparable with the uncorrelated case, they can be further analyzed on the basis of the two other quality metrics. Whitened ReciPSIICOS demonstrates noticeably lower radius of spreading at the expense of a slightly reduced detection ratio as compared to the ReciPSIICOS. We can also see that even in the synchronous case both proposed methods outperform the MNE. Figure 9 demonstrates distribution of the localization bias and point-spreading area for synchronous and asynchronous cases corresponding to SNR = 4 slice of dependencies in 8.A.

**Figure 9:**
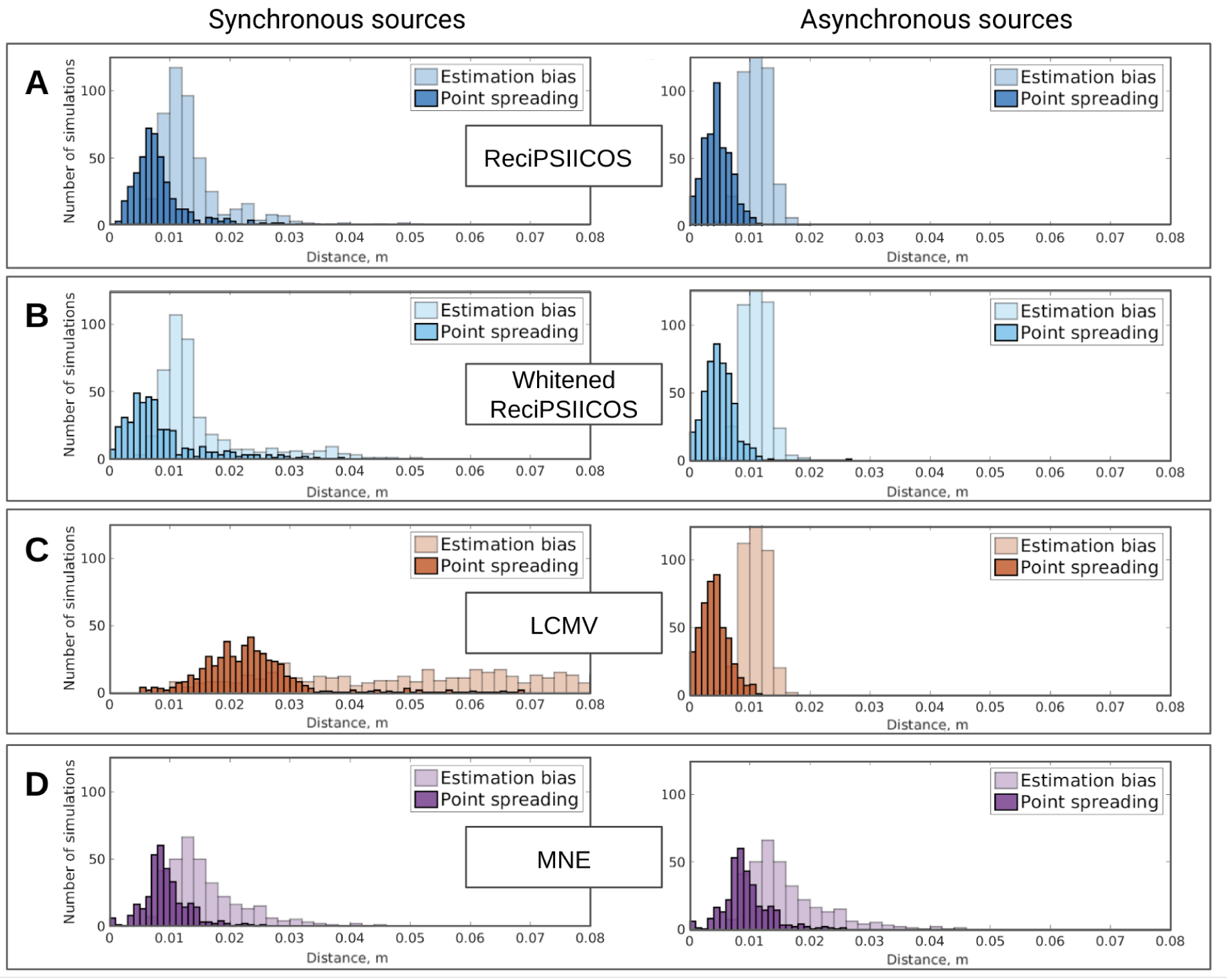
Distribution of localization bias and area of point spreading metrics for 500 Monte Carlo simulations. Noise level is adjusted so that SNR = 4. The results are computed for ReciPSIICOS (A), Whitened ReciPSIICOS (B), LCMV (C) and MNE (D).

Since the performance of all inverse solvers, and those based on the LCMV principle in particular, depends on the forward model inaccuracies, in panel B of Figure 8 we analyzed the extent to which the inaccuracies in forward modeling affected the three characteristics of the inverse solutions obtained by the four different solvers. For uncorrelated sources, one can observe that the localization bias is low enough for all methods, though it is slightly higher for MNE, and increases insignificantly with the increase of forward model error. As expected, the area of activity spreading grows for all beamformer-based methods, but remains within the acceptable range for the typical MEG forward model noise levels of 10% (Mosher et al. (1999)). MNE’s performance appears to be not adversely affected by the explored range of FM inaccuracy levels.

For correlated sources, the performance of ReciPSIICOS pars that of the MNE with a slightly higher detection ratio pertinent to the ReciPSIICOS. The Whitened ReciPSIICOS shows more compact solutions than the other three techniques and a higher detection ratio. In the synchronous case, due to general deterioration of performance of all four methods, the effects of forward model inaccuracies are significantly less pronounced than in the situation with asynchronous sources.

Projection rank is the only user defined parameter of the ReciPSIICOS and Whitened ReciPSIICOS techniques. Figure 10 depicts the three performance metrics considered above as functions of the projection rank for the two methods in synchronous and asynchronous cases. From these plots, both ReciPSIICOS approaches are characterized by a smooth and relatively flat performance profiles as a function of projection rank. Note that the projection rank has a different meaning for ReciPSIICOS and Whitened ReciPSIICOS. Increased projection for ReciPSIICOS corresponds to a less restrictive situation when the variance from the correlation subspace is expected to leak more intensively into the power-only subspace. At the same time, the greater the projection rank for the Whitened ReciPSIICOS method, the stronger is the suppression of the undesired variance from the correlation subspace, which happens at an expense of the variance in the power-only subspace. These simulations were conducted for the Neuromag probe and agree well with subspace power ratio plots in Figure 6.

**Figure 10:**
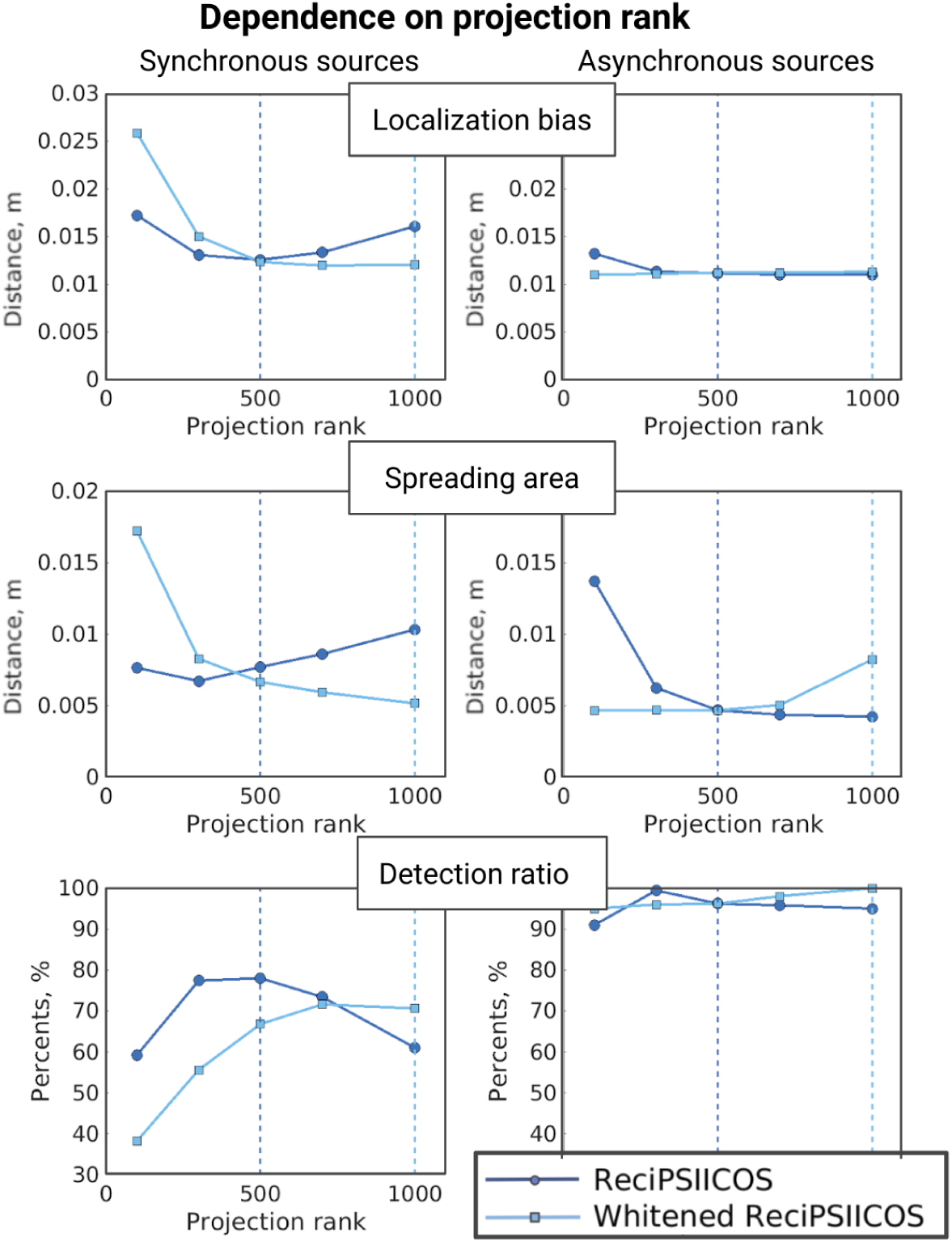
Dependence of localization bias, area of spreading and detection ratio on the projection rank for ReciPSIICOS and Whitened ReciPSIICOS techniques. Dashed lines shows the ranks picked for ReciPSIICOS and Whitened ReciPSIICOS analysis correspondingly.

#### 3.1.2 Three interacting sources

The majority of the present solutions to the correlated sources problem in beamforming (see Introduction) handle pairs of correlated sources by suppressing the source correlated with the target one by means of an extra zero constraint. In principle, these existing techniques can be extended to the triplets, quadroplets and etc. of potentially correlated sources. However, such an extension would lead to significant computational demands and would require methods for fusing the obtained results. The approach proposed here takes care of all correlated sources simultaneously. In this section we will focus on the example of three correlated sources. The data was modeled as described in section 2.6 and the three target sources were picked randomly, but so that they were no closer than 4 cm apart. We first consider sources activated with sinusoidal functions with a zero phase shift, but slightly jittered in time (see Figure 12.B). A representative case (a single Monte Carlo trial) is shown in Figure 11. Panel A illustrates locations of three randomly picked sources in the right frontal, right ventral and left parietal areas. We can see that in this scenario the LCMV beamformer produces completely incorrect activation map, ReciPSIICOS and MNE detect all three sources and demonstrate the comparable results. The Whitened ReciPSIICOS shows an outstanding performance, as the obtained activations are extremely focal and they successfully capture the initial activity.

**Figure 11:**
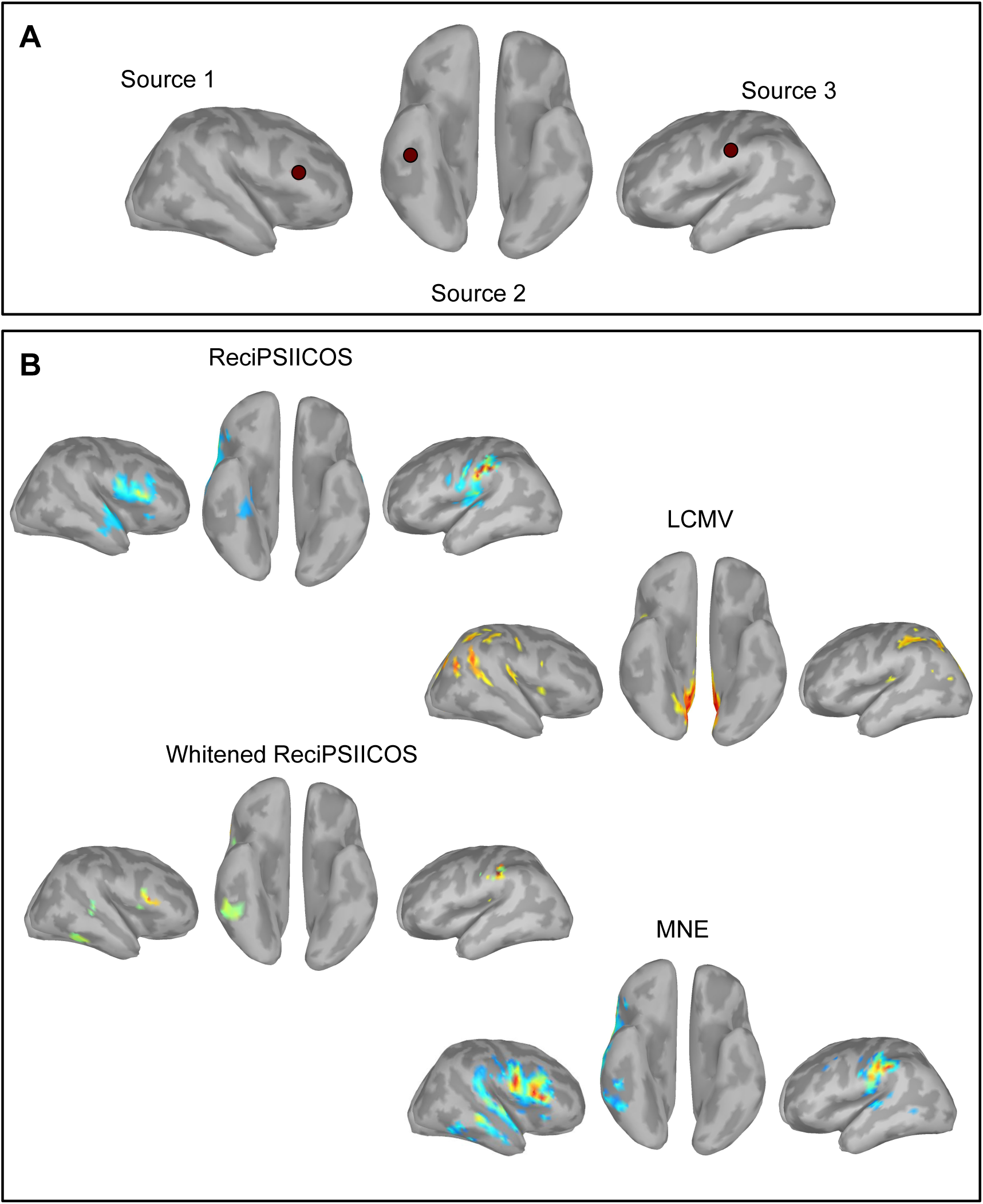
Modeling the case of three interacting sources. **A**. The locations of three randomly picked sources. **B**. Source activation maps reconstructed with ReciPSIICOS, Whitened ReciPSIICOS, LCMV and MNE.

**Figure 12:**
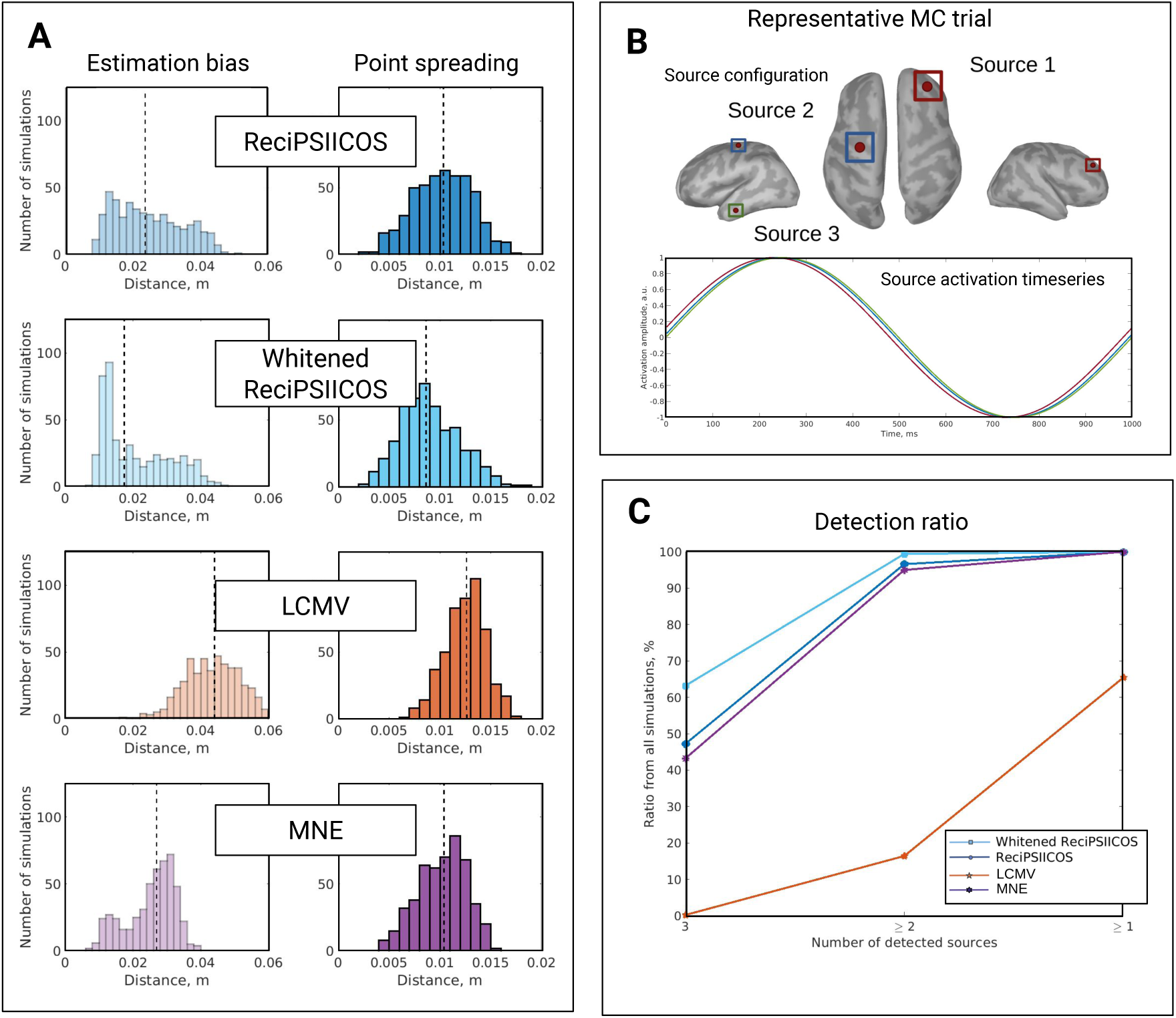
Simulation of three synchronously active sources. **A**. Distribution of two reconstruction quality metrics: estimation bias and point spreading value, for four reconstruction techniques: ReciPSIICOS, Whitened ReciPSIICOS, LCMV and MNE. **B**. One representative Monte Carlo trial: three cortical sources are randomly picked and activated with the synchronous sinusoidal functions with a random time jitter. **C**. Distribution of the number of detected sources for all simulations.

These findings are generalized using 500 Monte Carlo trials in Figure 12. Distribution of localization bias and point spreading radius for different methods over 500 Monte Carlo trials are shown as histograms in Figure 12.A. Dashed lines depict the median values. According to the first two metrics, it can be concluded that in the case of multiple active sources Whitened ReciPSIICOS significantly outperforms the competing methods, even the simple ReciPSIICOS.

#### 3.1.2 Three interacting sources

The majority of the present solutions to the correlated sources problem in beamforming (see Introduction) handle pairs of correlated sources by suppressing the source correlated with the target one by means of an extra zero constraint. In principle, these existing techniques can be extended to the triplets, quadroplets and etc. of potentially correlated sources. However, such an extension would lead to significant computational demands and would require methods for fusing the obtained results. The approach proposed here takes care of all correlated sources simultaneously. In this section we will focus on the example of three correlated sources. The data was modeled as described in section 2.6 and the three target sources were picked randomly, but so that they were no closer than 4 cm apart. We first consider sources activated with sinusoidal functions with a zero phase shift, but slightly jittered in time (see Figure 12.B). A representative case (a single Monte Carlo trial) is shown in Figure 11. Panel A illustrates locations of three randomly picked sources in the right frontal, right ventral and left parietal areas. We can see that in this scenario the LCMV beamformer produces completely incorrect activation map, ReciPSIICOS and MNE detect all three sources and demonstrate the comparable results. The Whitened ReciPSIICOS shows an outstanding performance, as the obtained activations are extremely focal and they successfully capture the initial activity.

These findings are generalized using 500 Monte Carlo trials in Figure 12. Distribution of localization bias and point spreading radius for different methods over 500 Monte Carlo trials are shown as histograms in Figure 12.A. Dashed lines depict the median values. According to the first two metrics, it can be concluded that in the case of multiple active sources Whitened ReciPSIICOS significantly outperforms the competing methods, even the simple ReciPSIICOS. ReciPSIICOS and MNE show the similar performance, as we could see previously with two sources. Classical LCMV has the worst performance and basically fails.

Panel C of the Graph demonstrates the distribution of the number of detected sources, where the first point corresponds to the ratio of all trials when all three ground-truth sources were found. Thus, in 63% of all simulations, Whitened ReciPSIICOS was able to localize all three sources while ReciPSIICOS performance was at 48%, MNE had a slightly lower performance at 44% and LCMV did not find the three sources in any simulation.

Figure 13 demonstrates the same metrics for the three moderately correlated sources activated with sinusoidal functions with 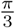 relative phase shift. This shift results in 0.5 pairwise correlation. As we can see, LCMV beamformer fails to detect three simulated sources in more than 80% of cases, whereas the methods from ReciPSIICOS family handle this situation well and loose one source only in 20% of cases. The bias and spreading metrics are also superior to the classical methods.

**Figure 13:**
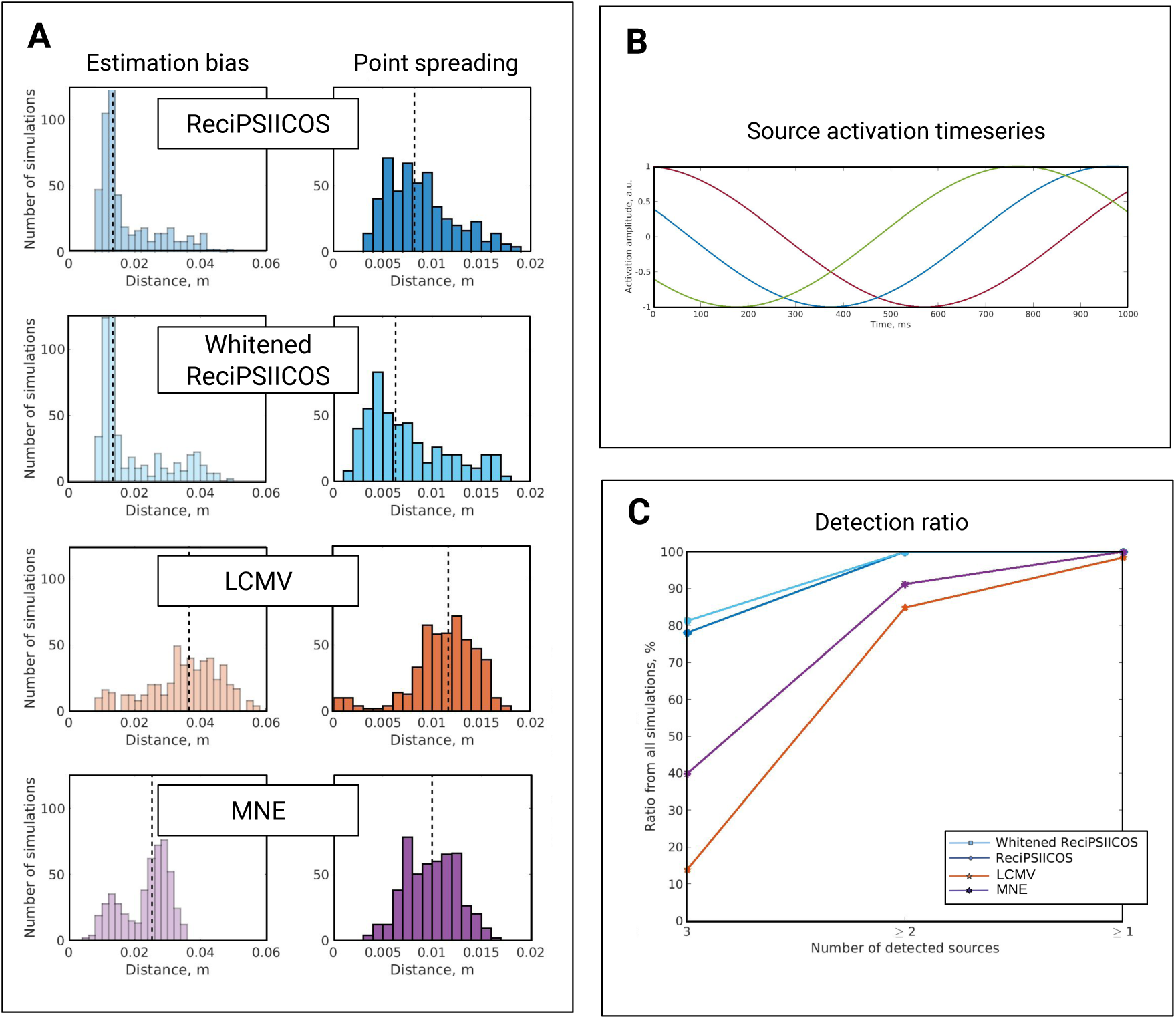
Simulation of three moderately correlated sources. **A**. Distribution of two reconstruction quality metrics: estimation bias and point spreading value, for four reconstruction techniques: ReciPSIICOS, Whitened ReciPSIICOS, LCMV and MNE. **B**. Example of representative trial: three cortical sources are randomly picked and activated with the sinusoidal functions with relative phase shift of 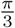 and random time jitter. **C**. Distribution of number of detected sources for all simulations.

### 3.2 Estimation of timeseries of correlated sources

Originally, beamformers were designed to estimate activity of a source at a given location. In this setting, when operating in the environment with correlated sources the LCMV beamformer will reduce the SNR of the correlated source timeseries estimates, see Section 2.2.2 for more details. This phenomenon is known as *signal cancellation*. When this happens the obtained timeseries will have a significant proportion of noise present in the beamformer timeseries estimates which will reduce the correlation between the estimated and true source timeseries. The proposed method for modification of the data covariance matrix allows for beamforming with significantly reduced signal cancellation and retained spatial selectivity properties.

To directly illustrate this we have performed 100 Monte-Carlo(MC) simulations where at each MC trial we simulated a pair of sources at random locations activated with harmonic timeseries submersed into the realistic brain noise, see Section 2.6 for a detailed description of the simulations. We performed six sets of 100 MC trials. Each set corresponded to different phase shift between the activation timeseries of simulated sources. We then used three different beamformers (LCMV and the two modified covariance beamformers), estimated the timeseries of the simulated sources and computed the correlation coefficient between the original and simulated timeseries for both sources. As we can see from both panels of Figure 14 the modified covariance beamforming scheme robustly assesses the shape of source activation timeseries for the entire range of the mutual phase lag values. At the same time, as expected, the adaptive LCMV beamformer achieves best performance only when the source timeseries are uncorrelated (Δ*ϕ* = *π*/2) and rapidly deteriorates as the phase difference decreases.

### 3.3 Real MEG data

In this section we describe the results of applying the proposed methodology to the real MEG datasets taken from two different experiments. The detailed descriptions of the paradigms and the datasets used are provided in section 2.8.

#### 3.3.1 Dataset 1

In this experiment, subjects received monaural auditory stimuli in the left ear. The following results were computed at a latency of 250 ms post-stimulus corresponding to the maximum amplitude of the response. Figures 15 and 16 show the estimated source power for both subjects and all considered methods. The source reconstruction revealed, as expected, most prominent activations in superior temporal gyrus, namely primary auditory cortex. Since the left ear received the stimulus, we expected to observe a stronger activation in the contralateral, right hemisphere and a weaker activation in the left hemisphere.

**Figure 14:**
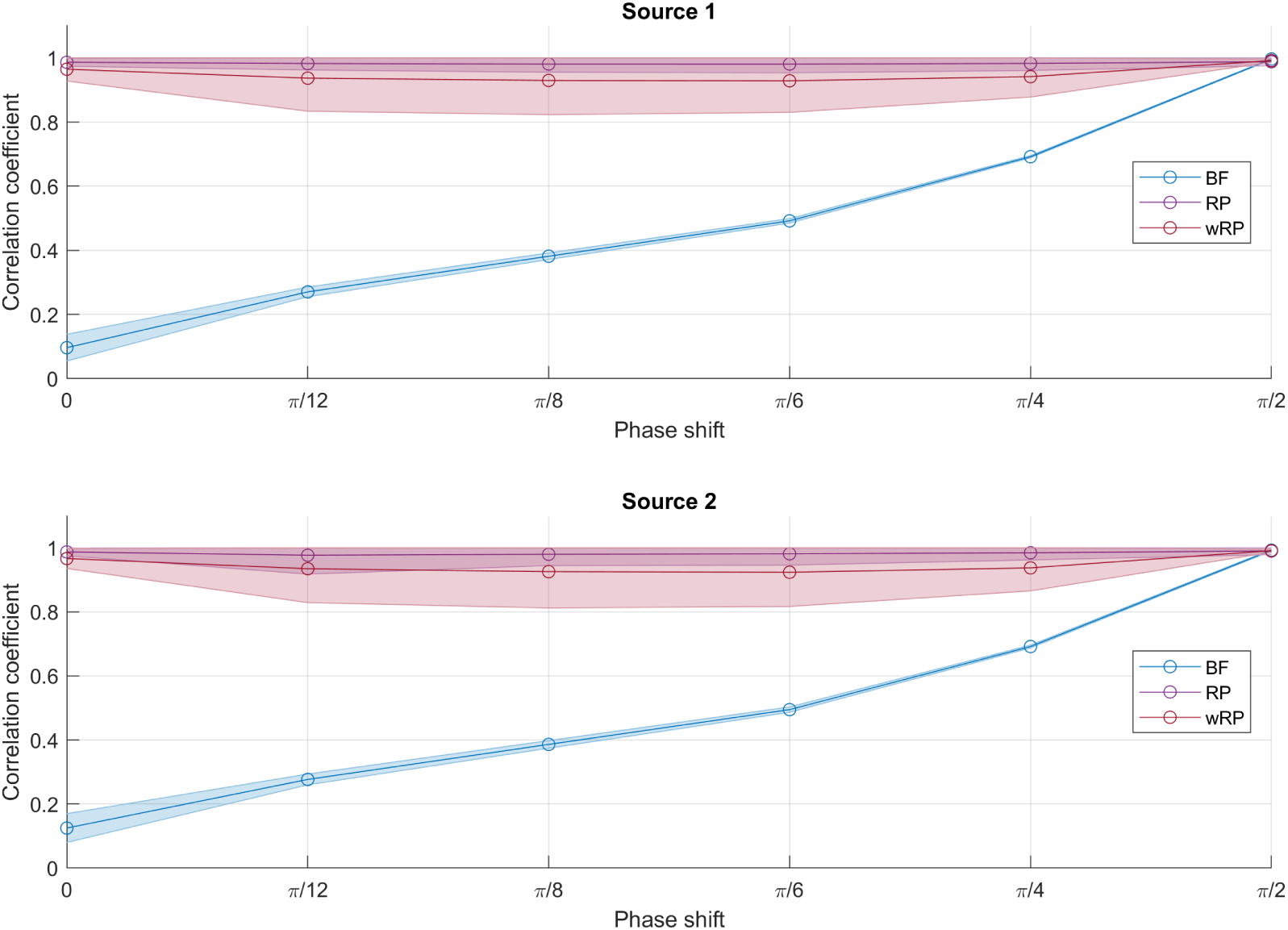
Pearson correlation coefficient between the original and estimated source timeseries for the three beamformers (LCMV, ReciPSIICOS and wReciPSIICOS) and six different phase lag values. The modified covariance beamforming scheme robustly assesses the shape of source activation timeseries for the entire range of the mutual phase lag values. At the same time, as expected, the adaptive LCMV beamformer achieves best performance only when the source timeseries are uncorrelated (Δ*ϕ* = *π*/2) and rapidly deteriorates as the phase difference decreases.

**Figure 15:**
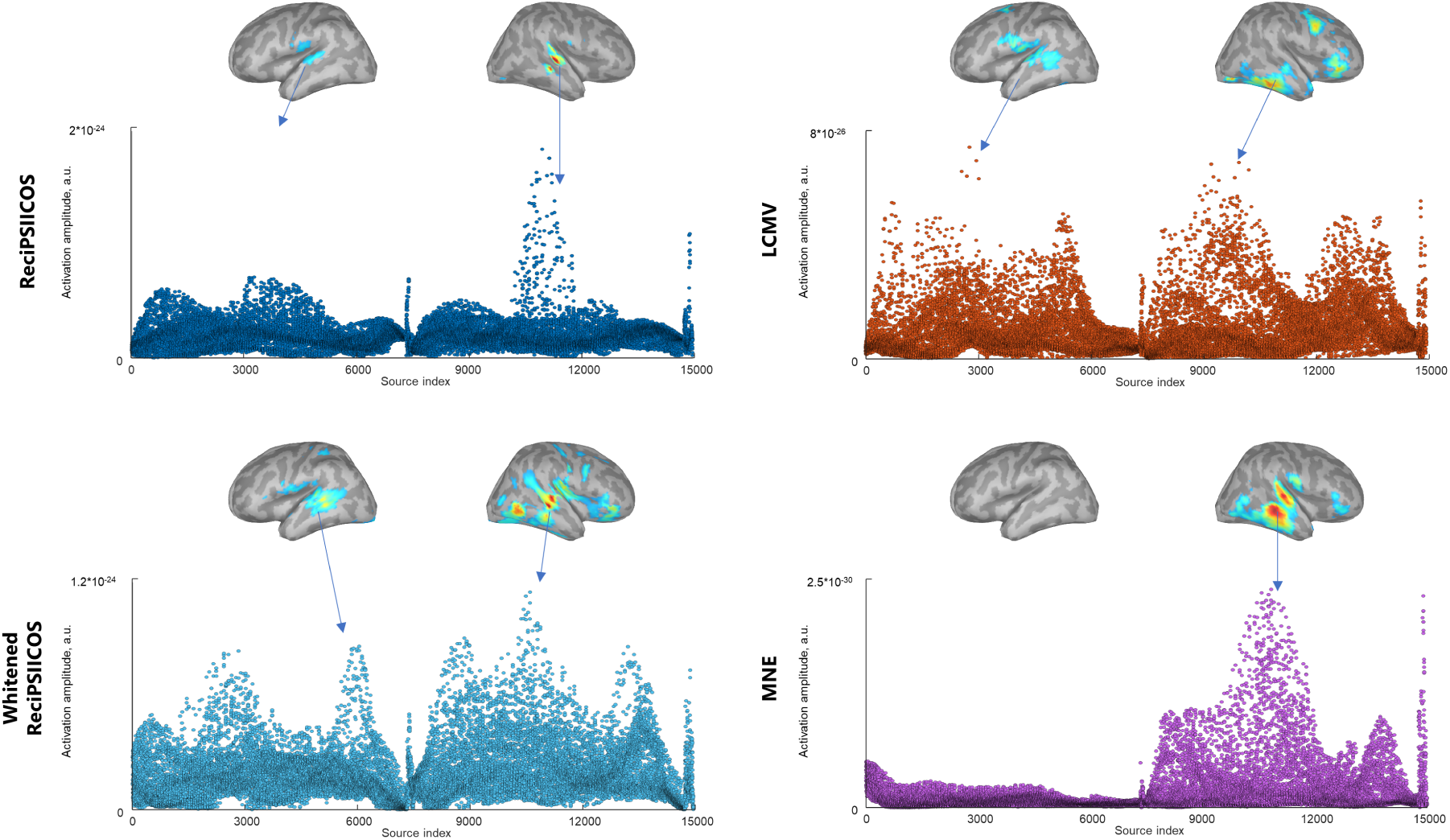
Power distribution of ASSR at 250 ms post-stimulus reconstructed with ReciPSIICOS, Whitened ReciPSIICOS, LCMV and MNE for the Subject 1.

For the first subject (Figure 15), it can be seen, that ReciPSIICOS, in comparison with LCMV beamformer and MNE, reveals more focal activations with the anticipated distinct maximum in the right auditory cortex and a lower, but still nonzero, activation in the left hemisphere. LCMV beamformer demonstrates a similar ipsilateral activation but an absolutely incorrect contralateral pattern. MNE detects only the right hemisphere blob activation. For this subject, Whitened ReciPSIICOS shows the results similar to the simple ReciPSIICOS, but the latter provides more focal activations. Notably, the amplitudes reconstructed with the LCMV beamformer technique are approximately 400 times lower in the dynamic range than those reconstructed with ReciPSIICOS, with the maximal power of 8 * 10^*-*26^ and 2 * 10^*-*24^ respectively.

The results for the second subject (Figure 16) are similar to those for the first one. The best localization is demonstrated by ReciPSIICOS, while LCMV does not capture the activity in the contralateral hemisphere and MNE localizes two spreaded activity blobs.

Figures 17 and 18 show the comparison of reconstructed source timecources for ReciPSIICOS and LCMV for two subjects, correspondingly. Panel A shows the calculated auditory steady-state responses at 40 Hz. Panel B demonstrates the primary auditory cortex source location which was highly active in both ReciPSIICOS and LCMV solutions. The timecources of the picked source are shown on Panel C with their envelopes. The blue line corresponds to ReciPSIICOS, and the orange solid line corresponds to LCMV. As the amplitudes of LCMV activations are by the order of magnitude lower than for ReciPSIICOS, for the visualization purposes, we showed the rescaled LCMV with orange dashed line and highlighted area. What is noticeable here is that for both subjects the ReciPSIICOS reconstruction reveals that the analyzed gamma activity is modulated at a 5-Hz secondary frequency (i.e. theta rhythm), which agrees with the previous study Doesburg et al. (2012). At the same time, the timecourse extracted by LCMV appears to be modulated at 10 Hz (i.e. alpha rhythm). Panel D shows the distribution of ratio of norms for the LCMV and ReciPSIICOS derived spatial filters. The median value is slightly lower than 1, which confirms that the observed amplitude differences can not be explained by the difference in spatial filter norms. The obtained results are consistent across subjects.

**Figure 16:**
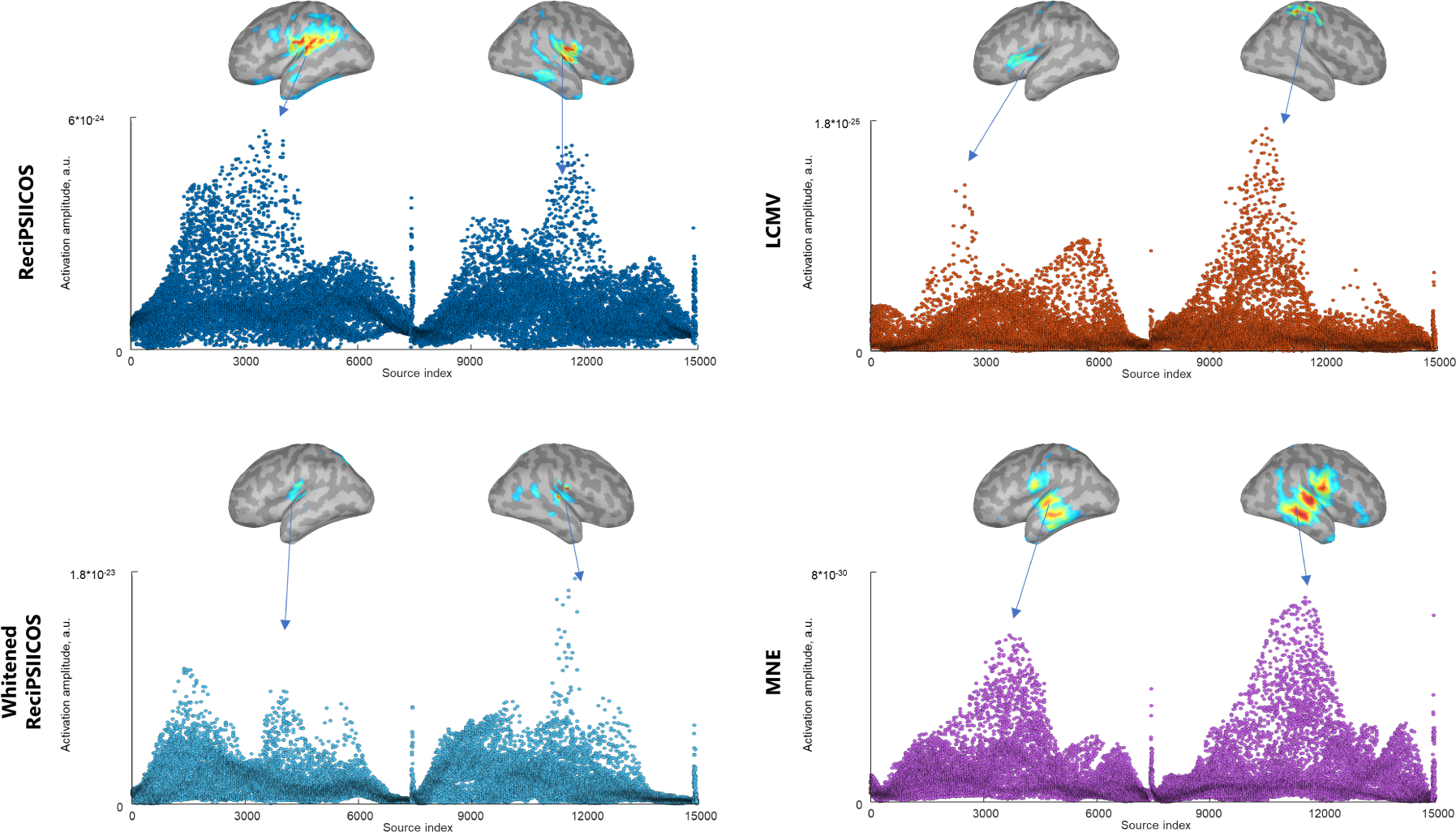
Power distribution of ASSR at 250 ms post-stimulus reconstructed with ReciPSIICOS, Whitened ReciPSIICOS, LCMV and MNE for the Subject 2.

**Figure 17:**
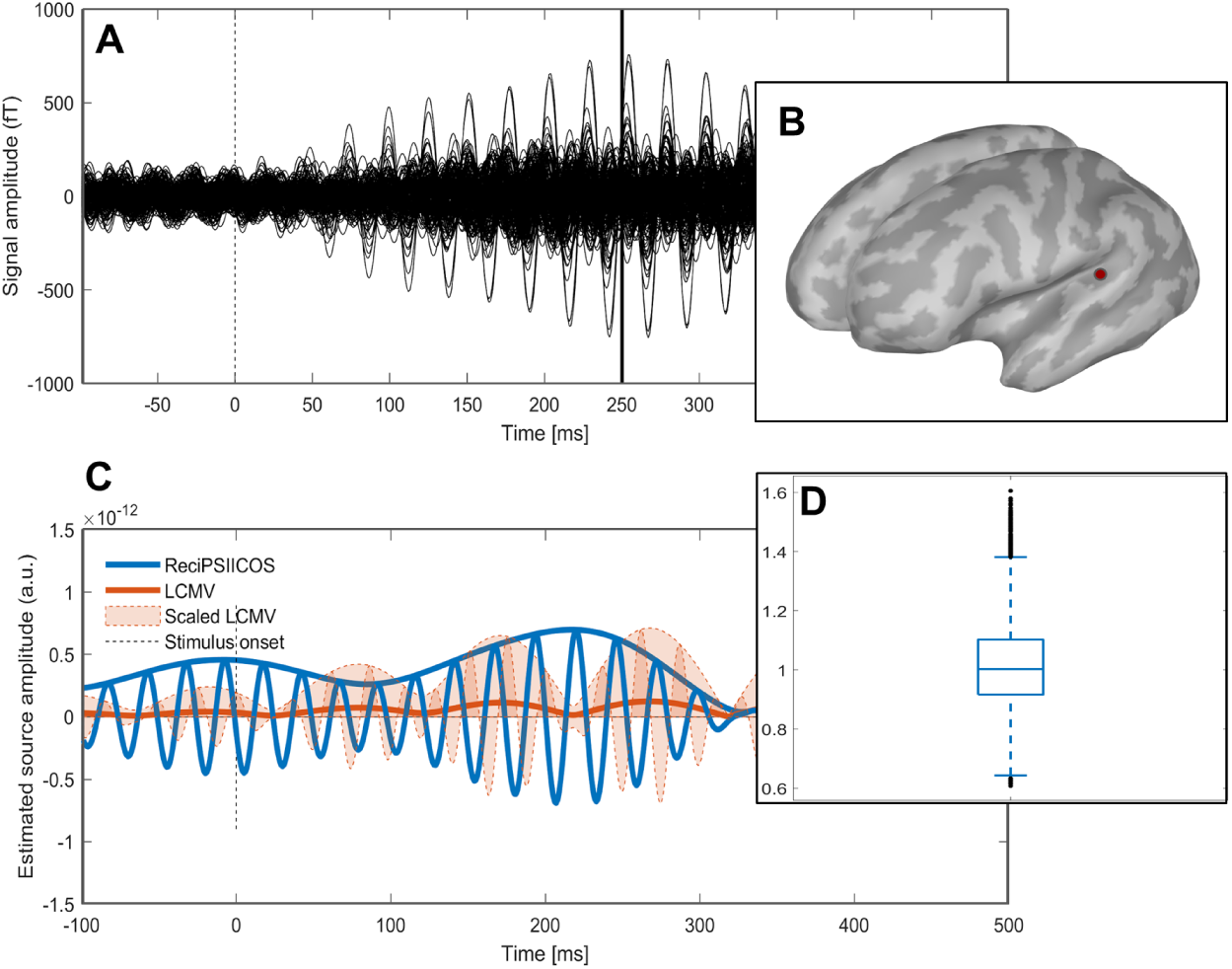
Analysis for Subject 1. **A**. Auditory steady-state responses at 40 Hz. **B**. The cortical source is highly active in both ReciPSIICOS and LCMV solutions at the gamma peak. **C**. The timeseries of a picked source in the primary auditory cortex reconstructed with ReciPSIICOS (solid blue line) and LCMV (solid orange line) techniques. ReciPSIICOS delivers the timeseries amplitude with the magnitudes that is significantly greater than that obtained with the original LCMV. To facilitate a meaningful comparison, an appropriately scaled timeseries estimate with LCMV technique is also shown. **D**. The distribution of ratios of LCMV weight norms to ReciPSIICOS weight norms calculated for each source.

**Figure 18:**
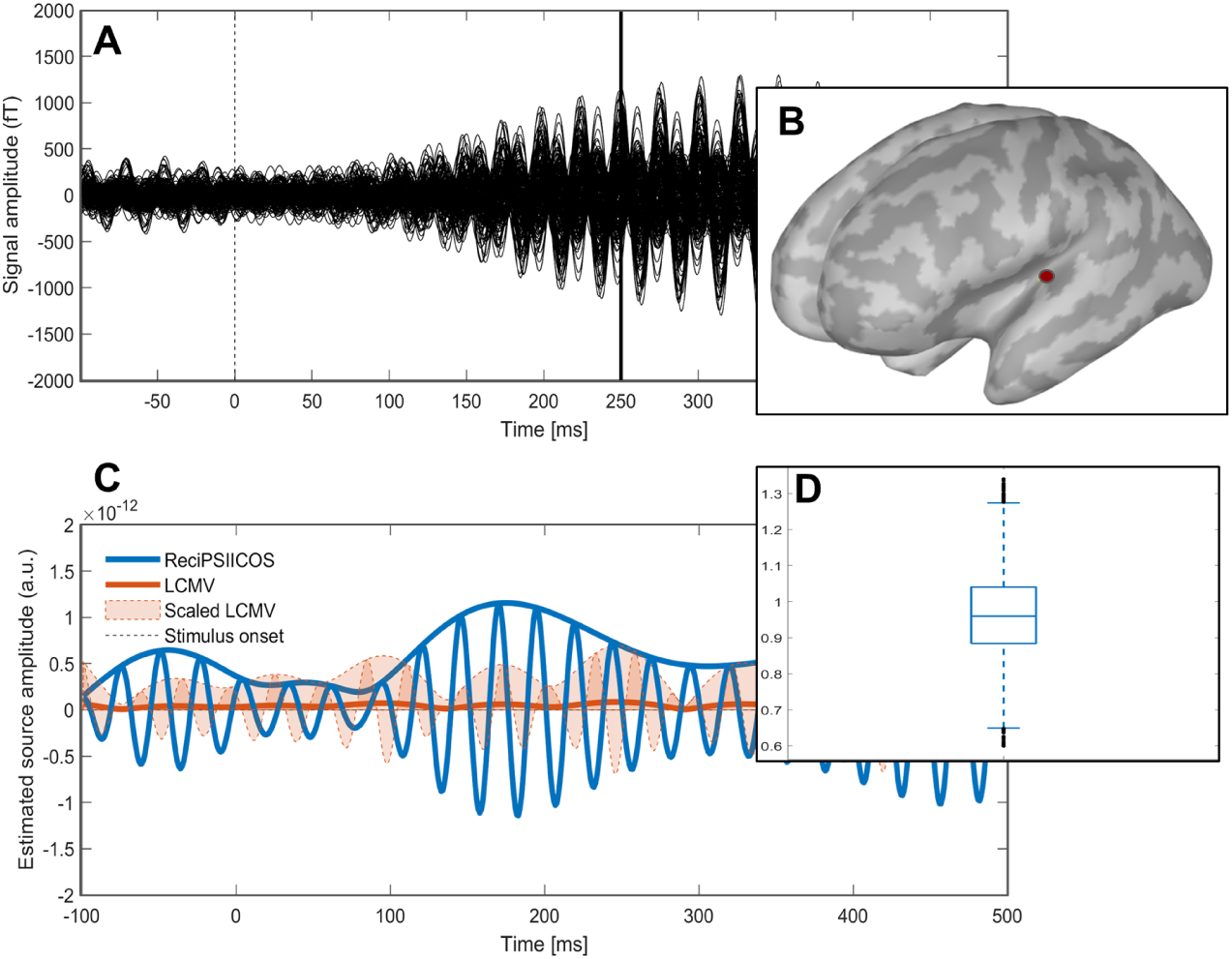
Analysis for Subject 2. **A**. Auditory steady-state responses at 40 Hz. **B**. The cortical source is highly active in both ReciPSIICOS and LCMV solutions at the gamma peak. **C**. Timeseries of the picked source in the primary auditory cortex reconstructed with ReciPSIICOS (solid blue line) and LCMV (solid orange line) techniques. ReciPSIICOS delivers the timeseries amplitude with the magnitudes significantly greater than that obtained with the original LCMV. To facilitate a meaningful comparison, an appropriately scaled timeseries estimate with LCMV technique is also shown. **D**. The distribution of ratios of LCMV weight norms to ReciPSIICOS weight norms calculated for each source.

#### 3.3.2 Dataset 2

The second dataset contains MEG data for the subject that participated in the auditory oddball paradigm and was instructed to press the button with the right index finger in response to the deviant stimuli. The stimulation was produced binaurally. Here we are focused on the localization of MMNm component (Näätänen et al. (1994)), so after the preprocessing described in 2.8.2 we calculated the differential ERF, which is shown on Figure 19, panel A. The peak of MMNm component is reached at 159 ms post-stimulus, so the inverse problem was solved for this time sample. Panel C shows the activation timeseries for one cortical source (Panel B) highly active both in ReciPSIICOS and LCMV solutions at the target time sample (159 ms post-stimulus). First, as expected, the amplitude reconstructed with LCMV (orange solid line) is significantly lower than the amplitude reconstructed with ReciPSIICOS (blue solid line). In order to compare these two solutions, we rescaled the LCMV solution so that the amplitudes at the target time sample were equal (the area highlighted in orange). It is clearly seen that ReciPSIICOS technique allows identifying the source that has one burst significantly different from the background activity. This burst corresponds to the MMNm component. The amplitude of the corresponding peak for LCMV beamformer is comparable with several other distributed along the whole timecourse. To prove that such a difference between the reconstructed amplitudes is due to the signal cancellation effect of the LCMV, but not due to the difference in the spatial filter norms, we calculated for each source the ratio of LCMV coefficients norm to the ReciPSIICOS coefficients norm. The obtained distributions are shown in Panel D. The mean value equals to 0.96 and the standard deviation is 0.14.

**Figure 19:**
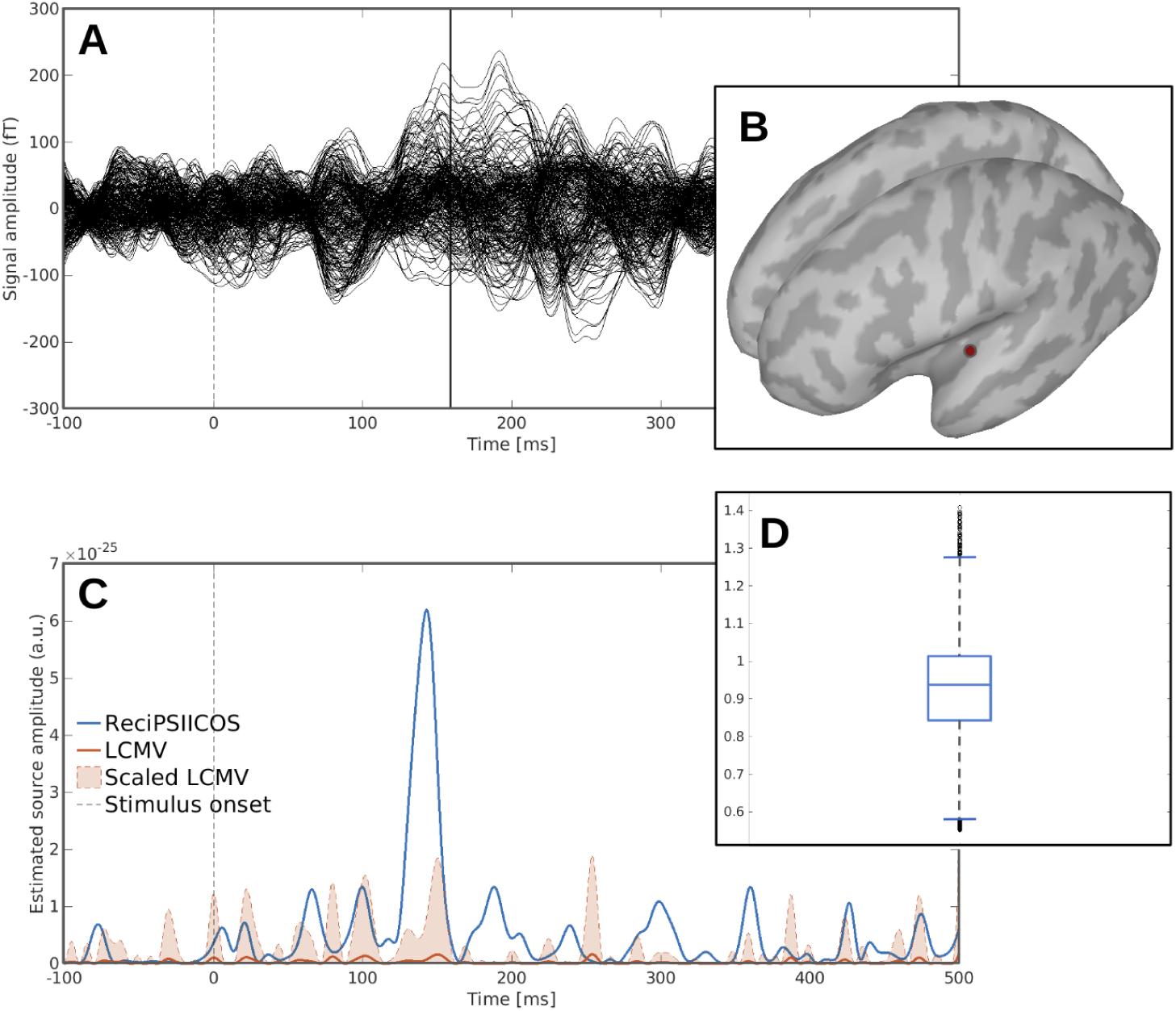
**A**. The differential ERF timecourses (deviant − standard responses). **B**. The cortical source highly active both in ReciPSIICOS and LCMV solutions at the MMNm peak. **C**. Timeseries of the picked source in the primary auditory cortex reconstructed with ReciPSIICOS (solid blue line) and LCMV (solid orange line) techniques. ReciPSIICOS delivers timeseries amplitude with magnitude significantly greater than that obtained with the original LCMV. To facilitate a meaningful comparison appropriately scaled timeseries estimate with LCMV technique is also shown. **D**. The distribution of ratios of LCMV weight norms to ReciPSIICOS weight norms calculated for each source.

Figure 20 demonstrates the estimated activation maps for ReciPSIICOS, Whitened ReciPSIICOS, LCMV and MNE. While the LCMV beamformer shows the high estimated amplitudes only in primary auditory cortex in the right hemisphere, ReciPSIICOS technique allows to localize activity in both hemispheres. At the same time, Whitened ReciPSIICOS technique shows similar activity in the primary auditory cortices in both hemispheres and an activation in the left motor cortex, which we expected to see due to the motor component of the task. MNE inverse solver found a highly spreaded activation and only in the left hemisphere.

**Figure 20:**
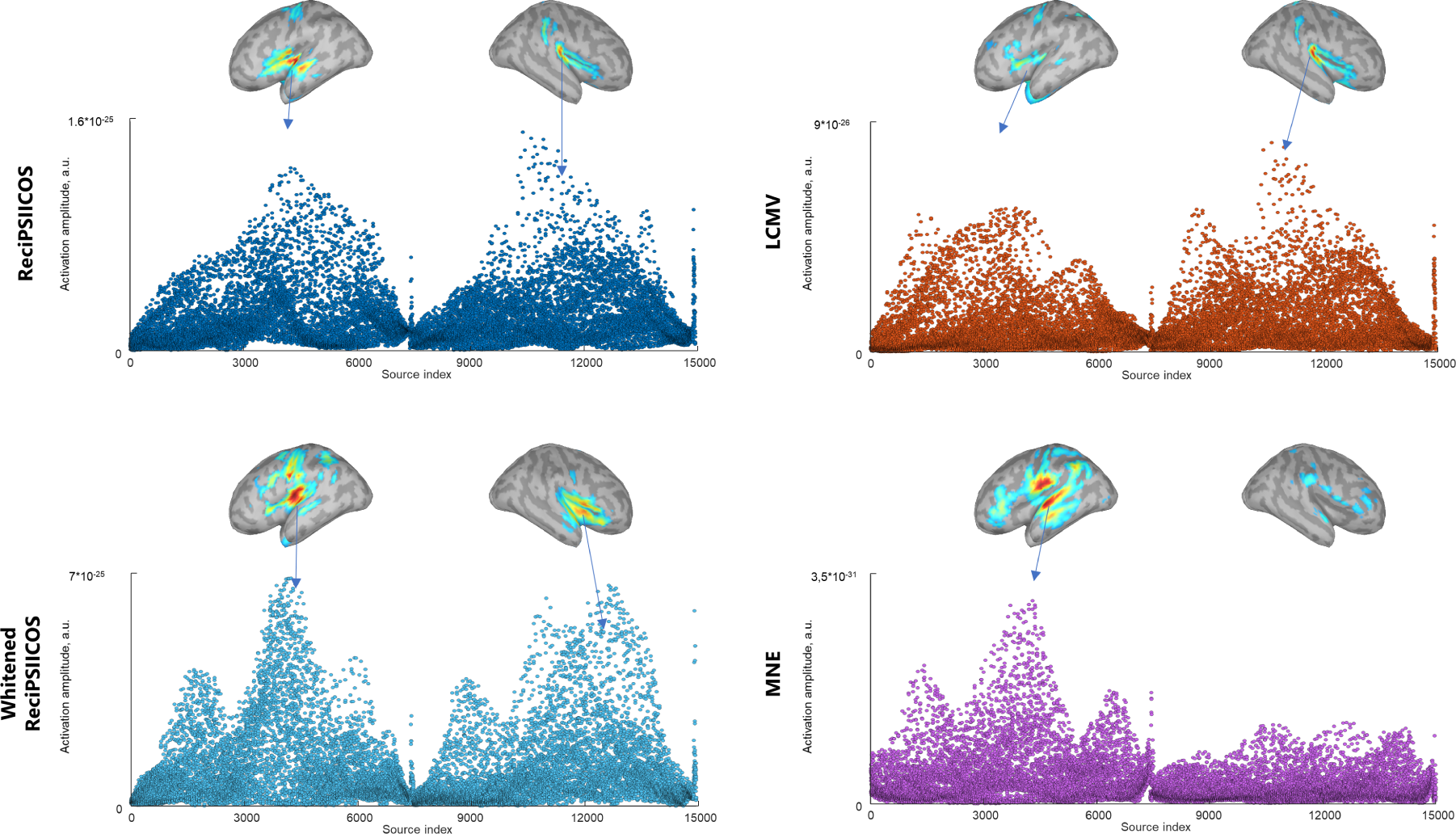
Reconstructed power distribution with ReciPSIICOS, Whitened ReciPSIICOS, LCMV and MNE for MMNm component at 159 ms post-stimulus.

Based on the above, we can state that the experimental data analysis results match those observed for simulated data. The performance of the LCMV beamformer appears to be significantly affected by the presence of correlations between cortical sources while ReciPSIICOS and Whitened ReciPSIICOS remain operable and deliver compact source maps with expected bilateral activation in the primary auditory cortex and a noticeably greater dynamic range.

#### 3.3.3 Non-positive definiteness induced by projection

The proposed approach uses a heuristics expressed in equation (25). This step is needed to guarantee that the data covariance matrix modified by the proposed projection remains positive definite (PD). The extent to which this manipulation is justified depends on the amount of energy stored in the negative eigenvalues of the reshaped and projected data. We assessed this amount using the expression (30) for the correlation matrices of real MEG datasets.

We estimated the contribution of negative eigenvalues into the total eigenvalue power in terms of *L*_1_-norm:

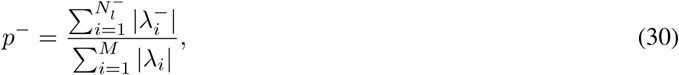

where *λ*_*i*_, *i* = 1 …*M* – all eigenvalues of the projected matrix 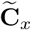 and 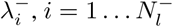 are the negative eigenvalues. The contribution of positive eigenvalues in the total power is then *p*^+^ = 1 − *p*^−^.

Figure 21 demonstrates the contribution of positive eigenvalues *p*^+^ as a function of projection rank for ReciPSIICOS and Whitened ReciPSIICOS calculated on Dataset 1 (panel A) and Dataset 2 (panel B). While ReciPSIICOS implies the projection on the power subspace, Whitened ReciPSIICOS performs the projection away from correlation subspace, so in order to align the results, each subplot on figure 21 has two x-axes. The lower x-axis corresponds to the ReciPSIICOS and its values are arranged in the descending order. The upper x-axis corresponds to the Whitened ReciPSIICOS and is arranged in the ascending order. Dashed lines show the ranks picked for the analysis. The ratio of positive eigenvalues differs from subject to subject, but in all cases, regardless of the MEG type (CTF or Neuromag), it reaches at least 80% and is typically higher.

**Figure 21:**
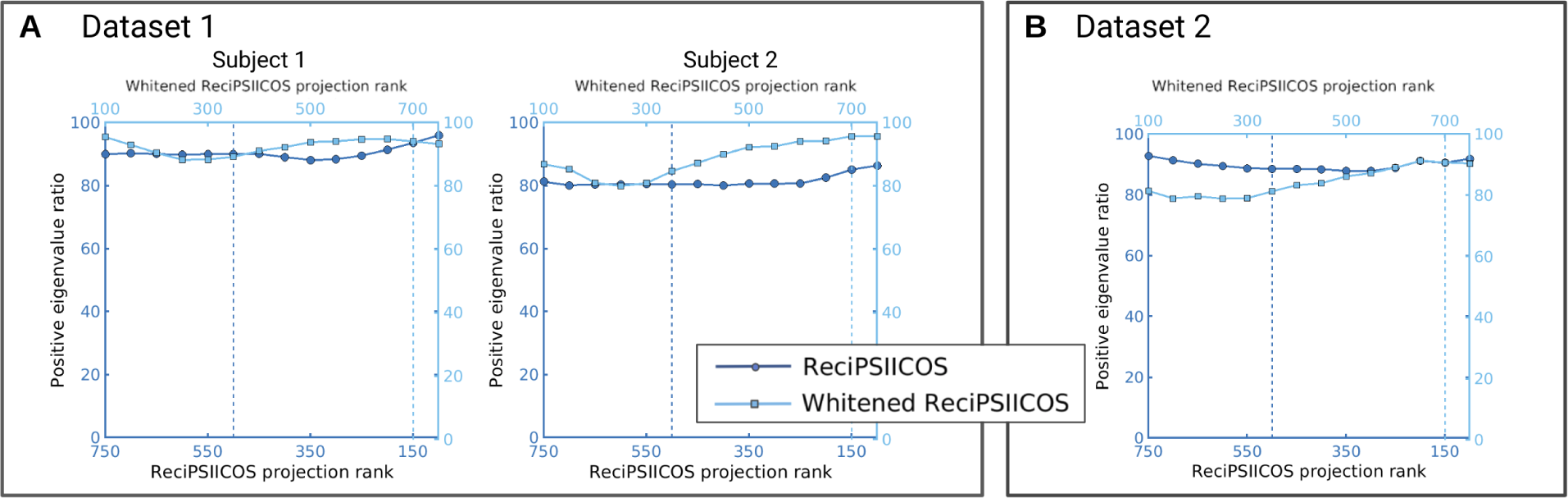
Contribution of positive eigenvalues of matrix 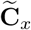 in total eigenvalue power as a function of the projection rank for ReciPSIICOS and Whitened ReciPSIICOS for the first MEG dataset (panel **A**) and the second dataset (panel **B**). Dashed lines show the projection ranks picked for the following analysis.

## 4 Discussion

In agreement with results of other groups and based on our own simulations, we can state that the adaptive beamformers outperform global methods for solving the inverse problem. However, the performance of adaptive spatial filters deteriorates significantly in the presence of correlated sources.

We have developed a novel method to supply robustness to the beamforming technique when operating in the environment with correlated sources. Our approach is based on the consideration of the sensor-space data covariance matrix as an element in a *M* ^2^-dimensional space. Using the MEG data generative model, we formulate the generative equation for the data covariance matrix and recognize that it contains contributions modulated by the diagonal elements of the source-space covariance matrix that span 𝒮_*pwr*_ subspace as well as its off-diagonal elements that represent coupling of neuronal sources and contributing variance to the coupling subspace 𝒮_*cor*_. Our method then builds a projector away from the subspace modulated by coupling and applies it to the data covariance matrix, effectively removing the contributions brought into the data covariance matrix by the non-orthogonality of the underlying source timeseries.

Strictly speaking, 𝒮_*pow*_ and 𝒮_*cor*_ subspaces overlap. However, the mutual spatial structure of the auto-terms and cross-terms allows us to partly disentangle the two subspaces and selectively suppress the variance in one of them while sparing the other.

We have developed two methods for building the projector. The first method simply fills *M* ^2^ × *N* matrix **G**_*pwr*_ with “topography vectors” as columns and attempts to find the subspace with the smallest dimension that captures maximum variance. This is done via SVD of **G**_*pwr*_ and the first *K* left singular vectors are then used to build the projector into the subspace they span.

The second approach is somewhat more complex. It projects the observed data correlation away from the 𝒮_*cor*_ subspace. To minimize the extent to which such a projection affects the variance in the 𝒮_*pow*_ subspace we perform the projection in the space whitened with respect to the spatial structure of 𝒮_*pow*_ subspace.

Both methods require specifying only a single parameter – rank of the projection *K*. In section 2.4, we suggest a natural procedure for the informed choice of *K* that is based on balancing the rate at which the variance is depleted from 𝒮_*pow*_ and 𝒮_*cor*_ subspaces with growing projection rank. Overall, we found that the method is quite robust (see Figure 10) to variations of this parameter in the broad range around the optimal value.

We have also shown that the proposed modification to the original beamforming approach is quite stable and stays functional when operated with realistically accurate forward models. Thus, the proposed technique does not require tuning many parameters and its simplicity is comparable to that of the MNE approach where only regularization parameter *λ* needs to be adjusted. Our simulations showed that for uncorrelated sources the proposed modification of the covariance matrix does not adversely affect the performance of the adaptive LCMV beamforming technique. Moreover, in the presence of sources with correlated activity the beamformer built on the basis of ReciPSIICOS projected covariance matrix remains operable and retains adequate localization performance unlike the classical LCMV beamformer. Therefore, the proposed approach can be considered as a universal tool for solving the inverse problem of MEG that exhibits super-resolution properties and remains functional in sub-optimal conditions.

Additionally, magnitudes of ReciPSIICOS beamformer weight vectors are comparable to those of the LCMV beam-former obtained using the original covariance matrix, as shown in Figure 19 (D). Nonetheless, the maximum output amplitude of the ReciPSIICOS beamformer exhibits a drastic growth (in some data up to two orders of magnitude!) as compared to the standard LCMV. This property allows for obtaining potentially more informative cortical activation maps of a greater contrast (Figures 15, 16, 20). We suggest that this result may indirectly gauge the prominence of source-space correlations typically present in the data.

Interestingly, the curves illustrating balance between the suppression of variance in the 𝒮_*pwr*_ and 𝒮_*cor*_ vary for two different MEG systems. According to the curves in Figure 6, sensor-array of the Elekta Neuromag system allows for a greater suppression of the undesired variance in the 𝒮_*cor*_ subspace practically without affecting that in 𝒮_*pwr*_ as compared to the probe of the CTF system. MEG systems based on the optically pumped magnetometers with the sensors located closer to the scalp capture higher spatial frequency harmonics of the magnetic field as compared to the traditional SQUID-based MEG machines. We suggest that the area under the curves shown in Figure 6 can be used as a design parameter to optimize the sensor layout of the future MEG systems.

As can be seen from Figures 19, 17, 18, the timeseries estimated with ReciPSIICOS beamformer are also characterised by a greater temporal specificty and greater SNR than those obtained with MNE or the original LCMV beamformer. The original LCMV clearly tends to cancel the two synchronous sources in the primary auditory cortex. Due to the comparatively low spatial resolution of the MNE, the obtained response combines activity of a large cortical area with potentially spatially varying response dynamics, which in turn results in low temporal specificity of the MNE-estimated timecourse.

The suggested projection procedure does not take into account the Riemannian nature of the manifold of the correlation matrices. Surprisingly, however, the eigenvalue spectrum of the ReciPSIICOS projected matrix remain largely positive. Only a small fraction of eigenvalues are negative and their magnitude is negligible and makes no more than 20% fraction of the total sum of the eigenvalue magnitudes. We return this projection back to the manifold of correlation matrices by simply replacing the negative eigenvalues with their absolute values. Although our results show that this is a reasonable working strategy, the proposed methodology would definitely benefit from the constraints imposed by the configuration of the natural manifold of correlation matrices. Approaches could be adopted in the future similar to the one described in Higham (2002) for finding the element on the manifold of correlation matrices that is the closest to the projected matrix. Overall, until a better solution is adopted to ensure the validity of the obtained imaging results, we recommend exploring the eigenvalue spectrum of the projected matrix and checking that the percentage of the “energy” brought into it by negative eigenvalues does not exceed the 10-20% threshold of the total sum of absolute eigenvalues.

Based on the simulations and experimental analysis, we conclude that the introduced here ReciPSIICOS and Whitened ReciPSIICOS procedures represent a simple, efficient and universal tool with super-resolution properties for solving EEG and MEG inverse problems in the environments with highly correlated neuronal sources. While the original adaptive LCMV beamformer performs suboptimally under these conditions, ReciPSIICOS family beamformers remain operable, retain super-resolution properties and maintain significantly higher detection ratio as compared to the MNE procedure known for its universal stability.

